# LeafRank: A phylodynamic framework for inferring relative fitness from single-cell phylogenies in chromosomally unstable tumors

**DOI:** 10.64898/2026.07.06.736651

**Authors:** Chenyu Wu, Kevin Leder, Zicheng Wang, Ruping Sun

**Affiliations:** Department of Laboratory Medicine and Pathology, University of Minnesota, Minneapolis, MN, USA; Masonic Cancer Center, University of Minnesota, Minneapolis, MN, USA; Department of Industrial and Systems Engineering, University of Minnesota, Minneapolis, MN, USA; School of Data Science, The Chinese University of Hong Kong (CUHK-Shenzhen), Shenzhen, China

## Abstract

Tumors contain cancer cells with diverse growth potentials that shape evolutionary trajectories, yet this fitness diversity remains difficult to quantify in cases of whole-genome duplication (WGD) and chromosomal instability. We present LeafRank, a mathematical framework that leverages single-cell DNA-seq phylogenies to infer the relative fitness of individual cells. Using a multi-type branching process model, LeafRank integrates full tree topology, including branch lengths and bifurcation patterns, to estimate marginal fitness probabilities under punctuated evolutionary regimes driven by rare driver events. To account for elevated aberration rates following WGD, we introduce a tree-rescaling strategy that adjusts for lineage-specific genomic instability. Unlike methods focused on predefined subclones, LeafRank ranks all sampled cells, enabling flexible assessment of growth heterogeneity. Simulations demonstrate high accuracy across spatial and non-spatial virtual tumors. Applied to ovarian cancer, LeafRank reveals directional and parallel selection in WGD tumors and identifies recurrent copy number events enriched in high-fitness lineages. WGD lineages do not show immediate growth advantages but acquire fitness through subsequent alterations.

## INTRODUCTION

Cancer evolution is the process by which cancer cells, expanding from a single founder cell, diverge by accumulating somatic aberrations, including single nucleotide variants (**SSNV**)^1,2^ and copy number alterations (**SCNA**)^3,4^, that can drive disease progression and resistance to treatment. The concept of somatic evolution, first articulated in Nowell’s seminal model of cancer evolution^5^, posits that subpopulations, or subclones, defined by distinct somatic aberrations within an established tumor can exhibit different survival and proliferation rates^6^, corresponding to differences in cellular fitness under a given microenvironment.

Fitness diversity among single cells within a patient tumor at the time of sampling reflects both cell-intrinsic growth potential driven by somatic variants and selective pressures imposed by the tumor microenvironment. Rapidly proliferating lineages indicate adaptive success at the time of sampling^7^, whereas slowly cycling cells may exhibit cancer stem-like properties^8–10^ or retain the potential to fuel treatment resistance^8,10,11^. Therefore, characterizing this fitness diversity is critical for addressing fundamental questions in cancer biology, including measuring selection pressures within the native tumor environment^12,13^, identifying somatic variants associated with fitness differences^14^, and predicting future adaptive potential^15^. However, in tumors with chromosomal instability (**CIN**), fitness diversity remains poorly characterized, and the fitness effects of SCNAs, including whole-genome duplication (**WGD**), remain elusive in patient tumor contexts.

Theoretically, subclonal fitness can be quantified by tracking the growth dynamics of cancer cell populations, ideally through longitudinal sampling during tumor evolution^16^. However, current technological constraints largely restrict such measurements to *in vitro* systems, mouse models^16,17^, or liquid tumors^18^. Consequently, despite the recognized importance of fitness diversity within solid tumors, methodologies that can infer subclonal fitness from patient tumor samples while preserving the native tumor context remain limited.

The relative fitness of a cell carrying an acquired somatic variant is commonly assumed to be reflected in the population size of the subclone harboring that variant, conditional on the time at which the variant arises^19^. Variant allele frequencies (**VAF**) of SSNVs measured from bulk sequencing have therefore been used as a surrogate for subclone size, providing evidence for detectable positive selection during subclonal evolution^12,20,21^. However, such analyses have been largely restricted to diploid genomic regions because SCNAs can confound VAF measurements, thereby excluding tumors with CIN phenotypes.

Notably, advances in single-cell DNA sequencing (scDNA-seq) have substantially improved the resolution of cell-to-cell genomic variation^22^, enabling the reconstruction of phylogenetic trees of detectable somatic variants in individual cells. Within such trees, subclone size can be estimated from the number of cells assigned to a given clade, while the birth time of a subclone can be approximated from branch lengths, or from the number of variants accumulated along the corresponding lineage, under appropriate molecular-clock assumptions. Both SSNVs and SCNAs have been used for phylogenetic reconstruction. For SSNVs, tree-sampling strategies have been developed^23,24^ to mitigate the impact of allelic dropout (ADO) in scDNA-seq data. In contrast, SCNAs can be robustly detected by leveraging read counts across large genomic regions, and the introduction of amplification-free scDNA-seq protocols^25,26^ has improved resolution of SCNAs to the megabase level. Despite the violation of infinite-sites assumption in SCNA evolution^27^, several methods use probabilistic frameworks to reconstruct SCNA histories. For example, MEDICC2 infers phylogenies by minimizing event distances between SCNA profiles and explicitly models WGD^28^, while SCICoNE^29^ and CONET^30^ jointly optimize tree con-struction and SCNA detection.

With phylogenetic trees derived from scDNA-seq data, mathematical modeling has begun to link information embedded in tree topology, including subclone sizes and branch lengths, to the population dynamics of cancer cells, enabling phylodynamic inference of subclonal fitness landscapes. SCIFIL infers fitness by modeling subclone sizes using a deterministic ordinary differential equation formulation of a branching process^31^. FiTree approximates subclone size distributions with a multi-type branching process under assumptions of large time and small mutation rates, and infers the fitness of specific genomic variants within a Bayesian frame-work that requires phylogenetic trees from multiple patients^32^. In contrast, cloneRate infers fitness by modeling subclone birth times or branch lengths using coalescent theory derived from a supercritical branching process^33^. SCPhyloX uses the Poisson distribution^34^ to model the expected branch length, including leaf-to-root and leaf-to-progenitor distances, to detect subclones under positive selection and estimate associated selection coefficients under as-sumptions of structured population growth^35^. Collectively, these methods rely on approxima-tions of branching process models, and focus on predefined subclones and typically assume uniform fitness within each subclone. Additional parameters, such as mutation rates, are also known to influence fitness inference^36,37^. Although joint estimation of multiple parameters has been proposed^38^, inferring absolute subclonal fitness levels remains challenging.

Rather than focusing on the absolute fitness values of specific subclones, we ask whether the relative fitness ranking of sampled single cells can be inferred from a phylogenetic tree reconstructed from genomic aberrations in a patient tumor. Leveraging a multi-type branch-ing process model, we developed **LeafRank**, a probabilistic algorithm that analyzes the full topology of phylogenetic trees derived from scDNA-seq data. Our approach incorporates es-tablished features of phylogenetic trees, including branch lengths and subclone sizes, while simultaneously accounting for additional unobserved factors that shape tree structure.

We designed LeafRank to accommodate the high CIN rates associated with WGD. LeafRank assumes branch lengths (aberration counts) are proportional to time. However, because base-line copy number (**CN**) dictates the substrate for subsequent alterations, ongoing CIN can al-ter aberration rates between lineages and violate this molecular clock assumption. Subclonal WGD is an extreme case that can significantly raise the baseline aberration rate^39–41^. To ad-dress this, we implemented a tree-rescaling approach that converts the input phylogeny into an ultrametric tree, allowing us to evaluate and correct for this molecular-clock violation.

This work is inspired by the non-cancer framework proposed by Neher et. al. which lever-ages influenza virus phylogenies to infer the relative fitness of viral strains under the assump-tion of small mutational fitness effects in a constant-sized population^19,42^. In contrast, we model cancer growth as an expanding population in which fitness differences between cell types can be large. Moreover, **LeafRank** does not assume the homogeneity of fitness within a subclone, and allows users to adjust the input parameters for the multi-type branching process model, thereby improving the flexibility in the evaluation of fitness diversity within the patient tumor. Using both non-spatial and spatial simulations^43^, we demonstrate that the inferred fitness rank-ings are generally robust to the choice of input parameters, with sensitivity arising only under specific parameter regimes. Applying **LeafRank** to copy number event trees from ovarian can-cer patient tumors undergoing WGD^44^, we provide support for fitness jumps between cell types, uncover evidence that post-WGD SCNAs reshape the fitness landscape, and examine the potential fitness consequences of WGD in patient tumors. For clarity, a guide to the terminology used throughout this paper is listed in Table S1.

## RESULTS

### LeafRank: A phylodynamic framework for inferring relative fitness from single cell trees

We developed **LeafRank**, a probabilistic framework for single cell phylogeny, inferring the relative fitness of sampled tumor cells. **LeafRank** takes as input a phylogenetic tree reconstructed from somatic aberrations, including SCNAs detected via scDNA-seq, alongside parameter configurations defined by a multi-type branching process model^45^ (Figure 1A). The framework out-puts the marginal probability of fitness types for leaf cells, conditioned on the observed tree topology and model configuration.

**Figure 1.**
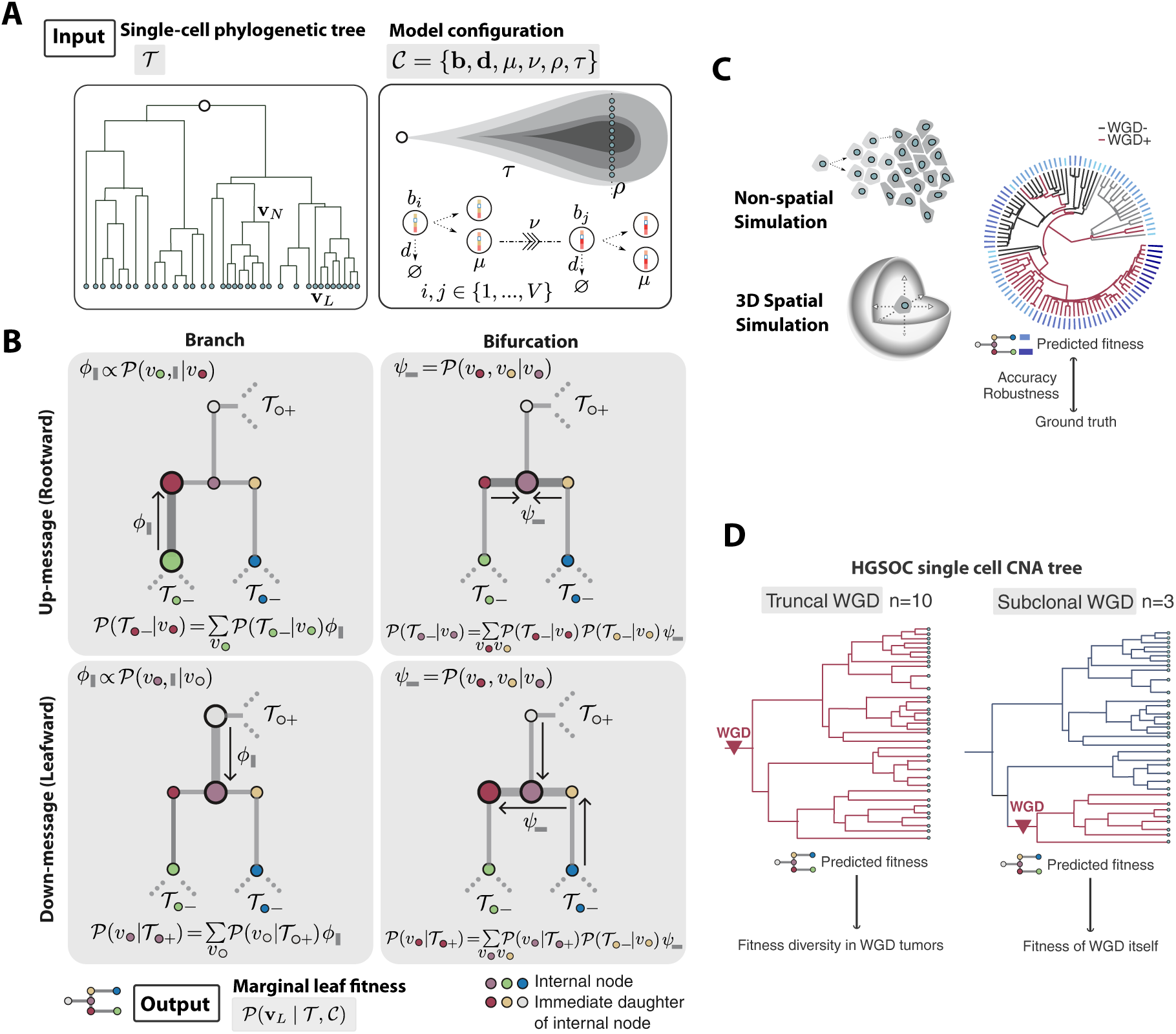
Study design and LeafRank framework. **(A)** The input for LeafRank consists of (1) a recon-structed single cell phylogenetic tree (binary bifurcation) derived from somatic aberrations (e.g., point mutations or copy number alterations) and (2) a multi-type branching process model parameter config-uration. Cancer cell expansion is governed by birth (**b**) and death (**d**) rates corresponding to one of V discrete fitness types. During division, daughter cells accumulate passenger aberrations at rate *µ* and may transition to higher fitness states at a driver aberration rate *ν*. In addition to the parameters govern-ing the branching process, the model incorporates a time-scale normalizing parameter *τ* and a single cell sampling probability *ρ*. The tree shape T integrates both branch lengths and bifurcation patterns, where **V**_L_ and **V**_N_ represent the fitness types of leaves and internal nodes, respectively. **(B)** Message pass-ing algorithm in LeafRank. To derive the marginal distribution of fitness types for sampled cells (leaves), LeafRank decomposes the probability calculation into the branch and bifurcation propagators. As shown in the row panels, the **“up-message” (rootward)** recursively collects the conditional probabilities of the subtree descending from each node. Conversely, the **“down-message” (leafward)** updates each node the marginal probability of its fitness types, conditioned on the observed phylogeny “above” the node, including all ancestral and sibling lineages. This bi-directional message passing facilitates the efficient computation of the marginal fitness distribution of all leaf cells. **(C)** Benchmarking via non-spatial and spatial simulations. Both non-spatial and spatial tumor growth models were employed to evaluate the accuracy and robustness of LeafRank, particularly in the presence of WGD. **(D)** Clinical application to HGSOC patient data. LeafRank was applied to HGSOC patient data to assess the fitness diversity of cells in tumors characterized by truncal WGD (Left) or subclonal WGD (Right).

Internally, **LeafRank** utilizes a multi-type branching process model that accommodates tran-sitions among discrete fitness states within an expanding population. This framework explicitly defines the probability density that an ancestral cell (an internal node), possessing a specific fitness type, generates the observed descending subtree (Figure 1B). Specifically, LeafRank decomposes this probability calculation into branch and bifurcation propagator functions. This formulation distinguishes our work from previous non-cancer frameworks^42^, which rely on con-tinuous diffusion approximations of small fitness effects. Crucially, this exact decomposition provides the foundation necessary to implement a message-passing algorithm, accelerating the computation of marginal fitness state distributions for each sampled cell (Figure S1). No-tably, the marginal fitness distribution of sampled cells takes into account the full information embedded in the tree topology, while the multi-type branching process framework naturally ac-counts for the stochastic nature of the clonal expansion dynamics. Detailed implementation of the **LeafRank** algorithm is provided in the Methods and Supplemental Text.

### Topological signatures of clonal expansion inform fitness ranking predictions

To verify LeafRank’s performance on ideal data, we first tested it on the ground-truth phylo-genetic trees and parameter configurations derived from virtual tumors whose growth satisfies our underlying assumptions (See Methods). Because LeafRank assumes branch lengths rep-resent elapsed evolutionary time, the input tree should be approximately ultrametric; that is, it should exhibit relatively consistent leaf-to-root distances when the samples originate from a single time point. We simulated virtual tumors using the multi-type branching process model, and randomly sampled single cells once the tumor reached a million cells. By recording the birth and death times of each cell during the simulation, we are able to reconstruct an accurate input tree, termed as a true time ultrametric tree or TT tree, as demonstrated in Figure 2. The simulation setting is detailed in Methods.

**Figure 2.**
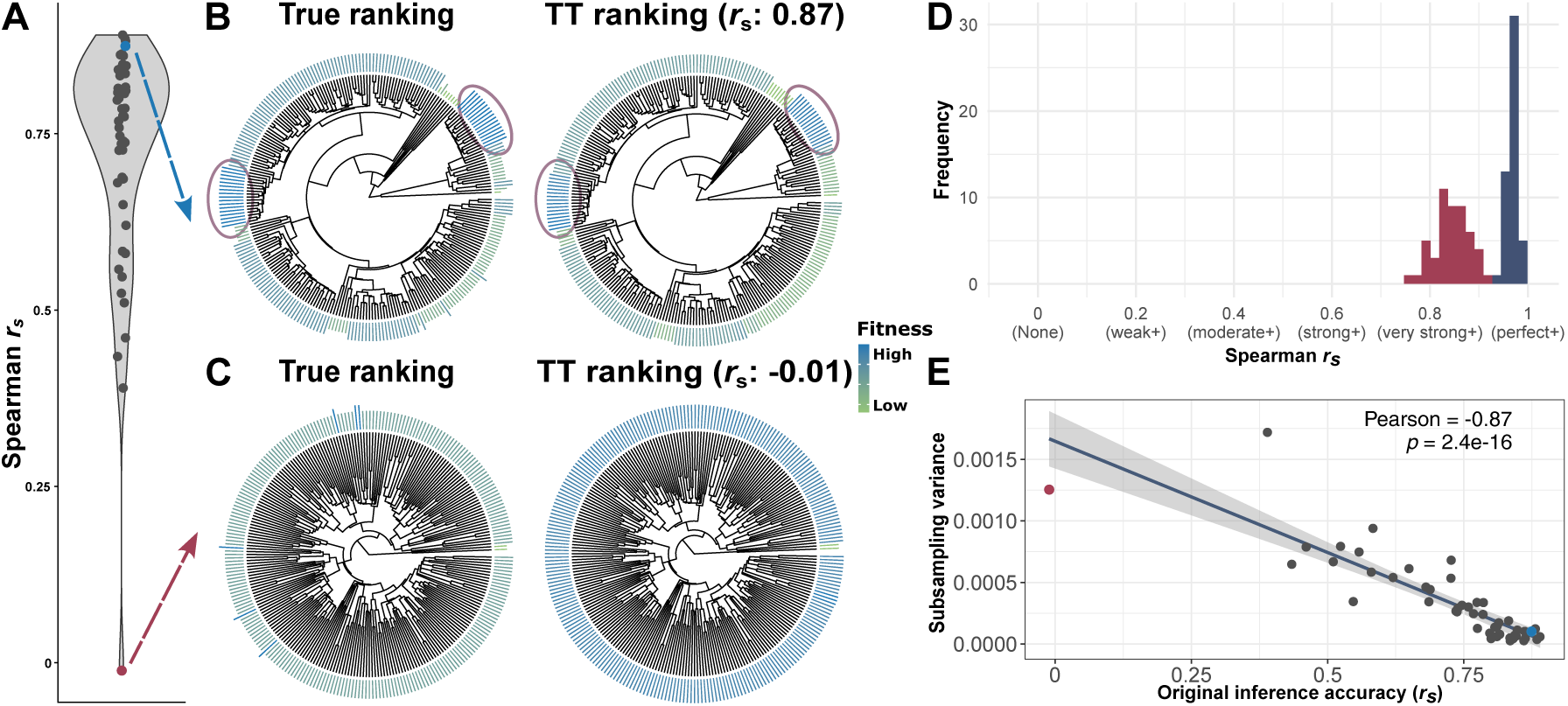
Phylogenetic tree topology enables robust inference of cellular fitness. **(A)** LeafRank performance across 50 simulated tumors, quantified by Spearman correlation between true and inferred fitness rankings derived from true-time (TT) phylogenies. **(B)** Representative example with high concordance (r_s_ = 0.87). Inferred rankings (outer ring) closely match the ground truth, accurately identifying high-fitness lineages (circles). **(C)** Representative low-concordance case (r_s_ ≈ 0), illustrating reduced agreement between inferred and true rankings. **(D)** Ranking stability to subsampling, measured as Spearman correlation between rankings inferred from subsampled TT trees and the original full-tree ranking for the two virtual tumors in panels B and C, respectively. Histogram colors match the points in panel A. **(E)** Inference accuracy is inversely associated with subsampling variability (Pearson correlation). Variability is defined as the variance in Spearman correlations across subsampled trees. Highlighted points correspond to the two virtual tumors in panels B and C.

LeafRank recovered fitness rankings in virtual tumors with high fidelity. Predicted fitness values were highly correlated with the ground truth, where 35 of 50 tumors exhibited a rank correlation exceeding 0.7 (Figure 2A). This result indicates that LeafRank accurately translates tree topology into cellular level fitness diversity, effectively distinguishing high-fitness lineages from slower-growing lineages (circles, Figure 2B). Topologically, these high-fitness lineages are identifiable by a “late but rapid” branching pattern, which arise later in time but showing significant expansion relative to the total sampled population.

We observe that LeafRank’s sensitivity in identifying high-fitness lineages is tied to the realized size of the corresponding lineage. In cases where driver aberrations occur extremely late, the resulting sublineages may lack sufficient time to expand or reach the detection threshold for sampling. Consequently, LeafRank failed to identify these isolated, high-fitness cells when their topological signal was not yet reflected in the sampled tree. This is exemplified by the case where LeafRank predicted a uniform fitness landscape, reflecting a sampled population dominated by fitness-homogeneous cells (Figure 2C). Here, emerging high-fitness lineages had not achieved sufficient size to be distinguished from the background. Consequently, the absence of a measurable topological signal led to a low correlation with the ground-truth rankings.

In contrast, cell lineages that failed to achieve sufficient clonal expansion were ranked as low-fitness, regardless of their early emergence. While these clones may branch early in the phylogeny, their limited sampled clone size indicates a lack of competitive advantage in growth, allowing LeafRank to correctly distinguish them from high-fitness expansions. In all the virtual tumor samples, LeafRank correctly located the lower fitness cells.

Because true fitness is not observable in practice, we asked whether the accuracy of LeafRank predictions could be assessed through the stability of rankings obtained from trees built on randomly subsampled cells. We found that, although fitness estimates for subsampled cells are generally well correlated with rankings from the full tree, the strength of this correlation varies across virtual tumors. For example, in the fitness-homogeneous virtual tumor, where LeafRank failed to pinpoint late selective lineages (red in Figure 2D), ranking stability under subsampling was substantially lower than in cases where LeafRank achieved high accuracy (blue in Figure 2D). Across virtual tumors, the variance in Spearman correlations between subsampled-tree rankings and full-tree rankings was inversely associated with prediction accuracy (Figure 2E). Together, these results suggest that ranking stability under subsampling can serve as a proxy for inference accuracy.

### Robustness of LeafRank to parameter uncertainty and evolutionary model spec-ification

After establishing LeafRank’s performance under ideal conditions (with TT tree and accurate input parameters), we next assessed its resilience to parameter uncertainty. In practice, these parameters often vary by biological context and are characterized by significant estimation uncertainty. We therefore examined how LeafRank responds to variations in these inputs to ensure the robust translation of tree topology into fitness rankings. By perturbing each parameter across a biologically plausible range, we found that while individual fitness values were affected in diverse ways, the overall ranking inferred by LeafRank remained largely consistent. Furthermore, by characterizing the effect of input parameter tuning on the probability distribution of predicted fitness values, we provide guidelines for achieving an informative ranking through the evaluation of fitness distribution.

We performed a univariate sensitivity analysis by perturbing the input parameters, including sampling probability, overall time scale, number of fitness states, birth and death rates, and driver aberration rate, often by an order of magnitude above or below the baseline values (Table 1). Despite these significant deviations, the fitness rankings re-inferred across the 50 virtual tumors (introduced in Figure 2) maintained accuracy comparable to that achieved using the ground-truth parameters (Figure 3A). These results highlight the robustness of LeafRank to parameter uncertainty and its ability to preserve the relative fitness hierarchy even across substantial input variation.

**Figure 3.**
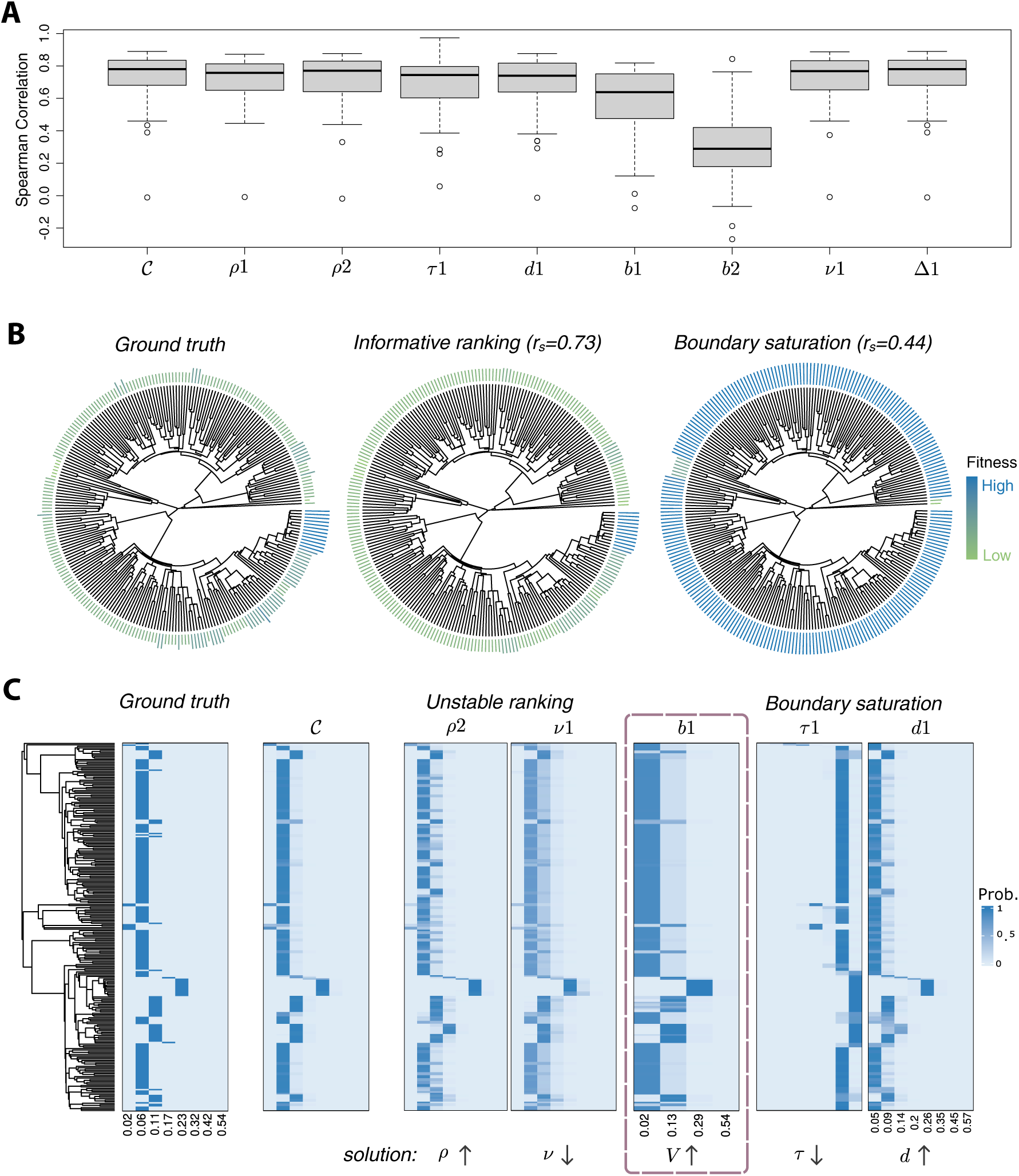
Perturbation analyses reveal robust rankings and guide stable input parameter calibration. **(A)** Robustness of LeafRank across eight input parameter perturbations, measured by Spearman correlation with the true fitness ranking (see Table 1). Boxplots summarize 50 simulated tumors per condition. **(B)** Example of reduced ranking resolution due to boundary saturation. Under the baseline parameter set (C), inferred rankings are well resolved, whereas increasing the time scale (*τ* = 8) compresses rankings toward the boundaries. Corresponding Spearman correlations (r_s_) are indicated. **(C)** Fitness-distribution heatmaps illustrating diagnostic patterns for tuning input parameter configuration to improve fitness inference. Rows correspond to individual cells and columns to fitness states. Two representative patterns are shown: unstable ranking and boundary saturation, each with two examples. Circled cases (b1) display both behaviors. Recommended parameter adjustments for each pattern are indicated below the corresponding heatmaps.

**Table 1.** Input parameter configurations for the univariate sensitivity analysis. Baseline values (*C*) correspond to the configuration used in Figure 2.

| ID | Parameters | Baseline Value ($\mathcal{C}$) | Perturbed Value |
| --- | --- | --- | --- |
| $\rho 1$ | $\rho$: Single cell sampling probability | 0.00025 | 0.001 |
| $\rho 2$ | | | 0.00005 |
| $\tau 1$ | $\tau$: Time scale normalizing parameter | 1 | 6 |
| d1 | $d_i$: Type i cell death rates | 0.18 | 0.15 |
| b1 | $b_i$: Type i cell division rates | $\{0.2 \cdot 1.2^i\}_{i=0,\dots,7}$ | $\{0.2 \cdot 1.53^i\}_{i=0,\dots,3}$ |
| b2 | | | $\{0.2 + 0.17 \cdot i\}_{i=0,\dots,3}$ |
| $\nu 1$ | $\nu$: Driver aberration rates | 0.0001 | 0.001 |
| $\Delta 1$ | $\Delta t$: ODE integration accuracy | 0.01 | 0.1 |

However, we observed that certain parameter configurations can result in uninformative fit-ness rankings that correlate poorly with the ground truth (Figure 3A). For instance, significantly reducing the number of fitness states (the b1 and b2 experiments) leads to a noticeable de-cline in accuracy. The b1 configuration performs better because it places greater resolution on lower-fitness phenotypes, consistent with the simulated distribution in which most samples have low fitness. By contrast, the b2 configuration assumes uniformly spaced fitness states across a predefined range, which does not capture this skew. Further details are provided in the Supplemental Text and Figure S2. These results suggest that while LeafRank is resilient to the scale of input parameters, it requires a sufficiently expressive model architecture to accurately resolve the fitness information. In light of these results we suggest using a V value of at least 7.

To provide practical guidelines for parameter optimization, we characterized how input settings shape the resulting fitness landscape, reconstructed from the inferred distributions across all sampled cells (Figure 3C). We observed two distribution patterns that emerge under spe-cific parameter regimes, revealing the conditions under which rankings become uninformative. These observations provide a framework for tuning LeafRank parameters to maximize the resolution and accuracy of the inferred fitness rankings.

The first pattern is ranking instability, characterized by closely related cells, such as those in sibling lineages, exhibiting sharply contrasting inferred fitness values. This pattern suggests that, under the current parameter regimes, LeafRank may be overly sensitive to stochastic noise embedded in the tree, leading to unstable predictions. For lineages with such erratic inferred fitness distributions, this excessive sensitivity to stochastic noise can distort fitness estimates at the individual leaf level, rendering the internal fitness structure biologically implausible. This instability typically arises when the cell sampling parameter *ρ* is set too small, the driver aberration rate *ν* is set too large, or the number of fitness types V is too small to resolve the underlying fitness structure.

The second pattern is boundary saturation, where a substantial fraction of lineages are inferred to have fitness at or near the preset minimum or maximum. This saturation reduces the resolution of the inference, obscuring the identification of high-fitness clones when most lineages are pushed to the maximum, or low-fitness clones when they are compressed at the minimum. This pattern suggests that the chosen parameters systematically deviate from the ground truth, forcing the inference to assign extreme values to reconcile the observed phylogenetic tree. Specifically, this arises when the time-scaling factor is either too small or too large, or when the birth/death rates are incongruent with the tree topology, such that tumor evolution along the given tree would require cells to have implausibly extreme fitness values.

Although each pattern may yield non-ideal fitness distributions, their individual effects on ranking accuracy, as measured by Spearman correlation, are minimal (Figure 3A). Empirically, boundary saturation reduced ranking accuracy more substantially than ranking instability as it flattens the tails of the distribution, obscuring differences between lineages with exceptionally high or low fitness. The most substantial deterioration in accuracy is observed primarily when both patterns occur simultaneously (Figure 3C, circled), highlighting the inherent robustness of LeafRank under unknown parameter settings. This robustness provides significant flexibility. For instance, if a user wants to focus on low-fitness lineages, they can increase the time scale *τ* to drive most lineages toward the maximum fitness state, thereby accentuating the contrast with the lowest-fitness lineage. Because the ranking remains stable, the lowest-fitness lineage can be consistently identified even when most lineages are concentrated at the highest fitness due to the chosen parameter configuration.

LeafRank provides curated parameter presets that users can adapt to achieve informative rankings (see Methods). Throughout the analyses in the manuscript, including applications to patient data, we implemented a distribution based quality control workflow to ensure that the resulting fitness distributions remained free of the two diagnostic patterns mentioned above.

### Robust fitness ranking in the presence of whole genome duplication and spatial sampling bias

To evaluate LeafRank under more realistic conditions, we tested its performance on trees containing noise and distortions typical of single-cell DNA sequencing data from tumor samples. While our numerical studies in previous sections used true time (TT) tree, empirical phylogenies are generally distance-matrix (DM) based, with branch lengths representing genomic divergence accumulated through events such as SSNVs and SCNAs. This discrepancy is often manageable under a constant molecular clock assumption^46,47^, whereby genomic divergence serves as a proxy for evolutionary time. However, in tumors exhibiting CIN, whole genome duplication (WGD) frequently occurs and violates this assumption. By doubling the DNA substrate, WGD accelerates the rate of subsequent genomic aberrations, thereby distorting branch lengths^39–41^. Therefore, genomic divergence in trees involving both WGD^−^ and WGD^+^ populations is no longer an appropriate proxy for evolutionary time. To account for aberration rate heterogeneity following WGD, we developed a modified *chronos*^46^ algorithm to recalibrate DM trees under a state-dependent molecular clock model that assigns distinct aberration rates to WGD^−^ and WGD^+^ branches (see Methods and Supplemental Text). The resulting WGD-adjusted ultrametric trees closely approximated the underlying TT trees.

When evaluating noisy reconstructed trees, WGD-adjusted trees substantially outperformed unadjusted DM trees in both non-spatial and spatial simulations (Figure 4F,G). Across all simulations, TT trees yielded the highest Spearman correlations with the true fitness ranking, providing an idealized upper-bound reference for inference accuracy. The WGD adjustment recovered rankings that closely matched this TT-tree benchmark. More specifically, the WGD adjustment corrected the spurious fitness inflation observed in WGD^−^ lineages. As shown in two representative virtual tumors (Figure 4B,C), DM trees systematically distorted the inferred fitness landscape. In these cases, the model misinterpreted the relatively shorter branches of WGD^−^ lineages as evidence of rapid expansion. By recalibrating branch lengths, the WGD-adjusted trees reduced this bias and restored correlations close to those obtained from the TT trees.

**Figure 4.**
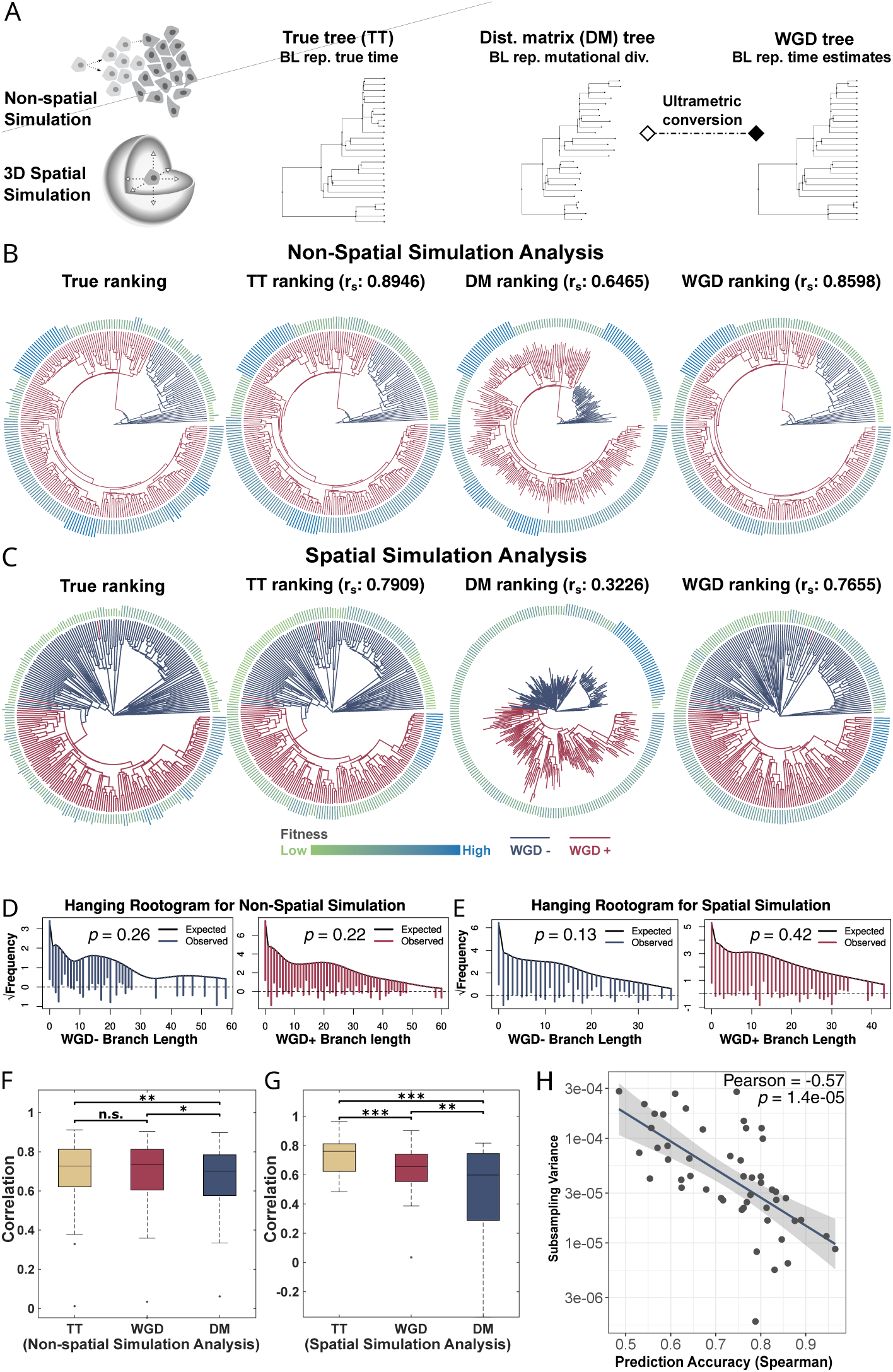
Benchmarking LeafRank in WGD-associated tumors using non-spatial and 3D spatial simulations. **(A)** *In silico* virtual tumor simulations generated under non-spatial and 3D spatial frameworks, each virtual tumor yields three phylogenies: a true-time (TT) tree, in which branch lengths represent evolutionary time; a distance-matrix (DM) tree reconstructed by neighbor joining, with branch lengths reflecting genomic divergence; and a WGD-adjusted ultrametric tree obtained by rescaling branch lengths as described in Methods. **(B,C)** Representative fitness inference from non-spatial **(B)** and spatial **(C)** simulations. True fitness rankings (outer ring, TT tree) are compared with rankings inferred from each tree type. Branch-level WGD status is color-coded according to the WGD status of sampled cells. **(D,E)** Hanging rootograms assess agreement between empirical branch-length distributions in DM trees and the expected frequencies predicted from WGD-adjusted trees. P-values were computed using a runs test of randomness on the positive and negative residuals along the branch lengths. **(F,G)** Inference accuracy across 50 virtual tumors per tree setting for non-spatial **(F)** and spatial **(G)** simulations, measured by Spearman correlation. Significance was assessed by Wilcoxon signed-rank test (n.s.: p ≥ 0.05, *: p < 0.05, **: p < 0.01, ***: p < 0.001). **(H)** Inference accuracy is inversely correlated with subsampling variability (Pearson correlation) in spatial simulations. Subsampling variability is measured similarly as in Figure 2E.

The branch length recalibration assumes that aberration counts follow a Poisson distribution. To evaluate this assumption, we used the temporal information from the WGD-adjusted ultrametric trees to calculate the expected Poisson-based branch length distribution, which represent the expected number of events per branch. We then compared it against the observed distribution from the unadjusted DM trees (see Methods and Supplemental Text for details). As shown in Figure 4D,E, the observed branch length distribution aligns closely with the distribution implied by the chronological time in the WGD-adjusted tree.

To evaluate LeafRank’s robustness against spatial structure-induced topological variations, we tested the framework on 3D virtual tumors^43^. Spatially structured populations result in sampling biases as cells from the same subclone tend to cluster in close proximity, which is commonly observed in solid tumors^48^. Indeed, this heightened complexity significantly degraded the performance of uncorrected DM-trees in the spatial simulation (Figure 4F,G). However, despite these multiple sources of noise, the WGD-adjusted tree significantly improved ranking accuracy. Notably, over 75% of these spatial virtual tumors achieved a Spearman correlation greater than 0.5, demonstrating the robustness of LeafRank in realistic evolutionary scenarios. By successfully restoring the expected relationship between branching density and evolutionary time, LeafRank is well-positioned for reliable fitness inference in solid tumors with CIN phenotypes.

Similar to the ranking stability test under subsampling shown in Figure 2E, we further exam-ine the relationship between ranking instability and accuracy in 3D virtual tumors (Figure 4H). Our analysis revealed that the negative correlation between the ranking accuracy and instability is preserved. This inverse relationship further suggests that subsampling-based ranking stability can serve as a valuable indicator of ranking accuracy in more realistic spatial simulated tumors, particularly relevant in patient tumors when the ground truth is unknown.

### Lineage fitness landscapes of ovarian cancer reveal ongoing selection following truncal WGD

We applied LeafRank to an scDNA-seq dataset of high-grade serous ovarian cancer (HGSOC) comprising treatment-naive samples collected from both primary and metastatic sites^44^ (see Methods). This cohort exhibits highly recurrent WGD characterized by both truncal and sub-clonal events, including cases of independent, parallel WGD alterations within a single patient’s tumor lineage. The recurrent yet variable distribution of WGD provided an ideal framework to investigate the fitness landscape of genome doubled cells and the specific selective advantages conferred by WGD.

We first investigated whether WGD, occurring as a truncal event, acts as a dominant driver to produce a uniform fitness landscape across the resulting population, or if subsequent outgrowth involves strong selective events occur post-WGD. To this end, we analyzed ten patients with truncal WGD^44^ whose samples were collected from the same anatomical site (including nine metastatic lesions, Figure S3 and Figure S4), minimizing sampling biases caused by spatial divergence. In these tumors, because WGD is a truncal event, we hypothesized that lineages arising after WGD evolve at a similar SCNA rate. Consequently, we converted DM trees reconstructed using MEDICC2^28^ into ultrametric trees using the constant rate *chronos* method (see Methods). LeafRank produced robust ranking inferences from both the original DM trees and the transformed ultrametric trees, while the rootogram test indicated that the constant rate model adequately captured SCNA accumulation in the majority of truncal WGD tumors (Figure S3 and Figure S4). For consistency, we present results based on the ultrametric trees. Details about the analysis are provided in the Supplemental Text.

LeafRank uncovered substantial fitness jumps during the tumor growth in over half of these patient tumors. In metastatic tumor OV-105 (sampled from omentum), for example, we defined 16 potential fitness types (0-15) with birth rates increasing by a multiplicative coefficient of 1.1. LeafRank revealed three major fitness groups, identified as types 6, 9 and 10, emerging through successive fitness changes toward the upper portion of the phylogenetic tree (Figure 5). This pattern resembles directional selection^49^, in which fitness rises as cells approach a specific aneuploidy state. Notably, these transitions varied in magnitudes. A major leapfrog occurred from the ancestral type 0 to type 6, followed by another significant jump to type 9, and a final incremental step to type 10. These results suggest that post-WGD evolution is characterized by ongoing selection rather than neutral expansion. The discrete fitness changes imply the presence of specific driver events coincident with lineage branching (Figure 5A). Furthermore, the heterogeneity in selective advantage confirms that a multi-type branching process is more appropriate than a continuous weak fitness-gain model^42^ for capturing inter-clonal disparities.

**Figure 5.**
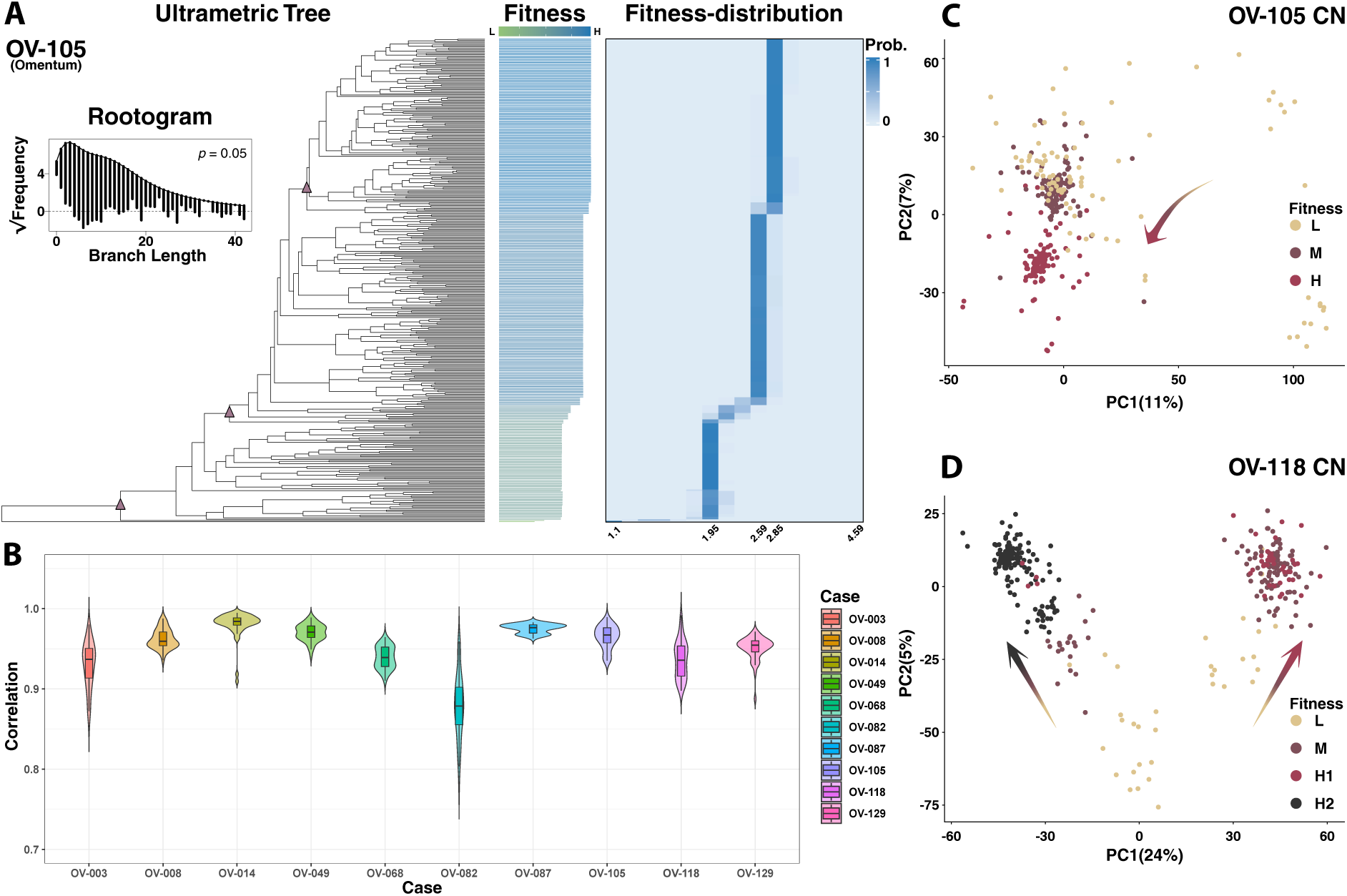
Detection of selection in HGSOC patient tumors with truncal WGD. **(A)** Representative patient tumor OV-105 showing directional selection in the single-cell phylogeny. Three subpanels are shown for OV-105 from left to right: (1) the Ultrametric Tree subpanel displays the rescaled tree de-rived from the copy-number (CN) event–based phylogenetic tree generated by MEDICC2, along with a rootogram visualizing the agreement between the empirical branch-length distribution in the MEDICC2 tree and the expected frequencies predicted from the rescaled tree. Three purple triangles mark parental nodes at which driver events are presumed to have occurred based on fitness predictions of sampled cells. (2) The Fitness subpanel reports the mean inferred fitness for each cell, as predicted by LeafRank. (3) The Fitness-distribution subpanel illustrates the probability distribution that each cell belongs to pre-defined cell types, with each cell type corresponding to a distinct fitness value. **(B)** Stability of fitness ranking assessed using leave-m-out subsampling across 10 truncal WGD HGSOC tumors. Violin plots show the distribution of Spearman correlations between fitness rankings inferred from the full tree and those obtained from 50 subsampled trees. **(C,D)** Patterns of directional and parallel selection reflected in CN evolution. Principal component analysis of genome-wide CN profiles is shown for patient tumors OV-105 **(C)** and OV-118 **(D)**. High-dimensional CN profiles from single cells are projected onto the first two principal components (PC1 and PC2), with the proportion of variance explained by each component indicated. Cells are color-coded by inferred fitness: low (L), medium (M), and high (H). Arrows indicate evolutionary trajectories in CN space underlying fitness changes. Fitness increases in OV-105 along a unidirectional trajectory (directional selection), whereas OV-118 exhibits two parallel trajectories (parallel selection).

In patient OV-105, the population sizes of the three major fitness groups were comparable, indicating that high-fitness cells have not yet dominated the population at the time of sampling. This observation highlights that sampled tumor cells represent a snapshot of an evolving population, where lineage fitness is not reflected by the current abundance. Specifically, of two lineages with identical sample sizes, a younger lineage possesses significantly higher fitness than an older one. By jointly accounting for both the relative size of a population and the age of the lineage, LeafRank distinguishes between lineages that are large simply because they had more time to grow and those expanding rapidly due to a true selective advantage.

To evaluate the reliability of LeafRank in these patient tumors where the ground truth is lacking, we utilized subsampling variability as an indicator of accuracy through a leave-m-out sub-sampling strategy. After estimating relative fitness values for all sampled cells in the original phylogenetic tree, we randomly subsampled 80% of the leaves and re-inferred fitness for each subsampled tree. This procedure was repeated 50 times, generating a distribution of inferred fitness rankings across replicates. As shown in Figure 5B, the subsampled trees consistently preserved high correlations (r_s_ ≈ 0.9) with the original rankings across all ten patients. These results demonstrate that LeafRank provides robust and consistent fitness inference under sub-sampling. The small subsampling variability observed in these patient tumors suggests high accuracy of LeafRank, consistent with the stability-accuracy relationship revealed in our simulation studies.

In addition to the directional selection observed in OV-105, we also identified a pattern of parallel selection, whereby multiple lineages with distinct CN profiles independently acquire fitness advantages (Figure S4). For example, in OV-105, the fitness progression from low to medium to high (L-M-H) follows a unidirectional trajectory in the CN landscape, as reflected by the principal component analysis (PCA) of single cell CN profiles (Figure 5C). In contrast, OV-118 exhibits two parallel trajectories in the CN space (Figure 5D). Here, a low fitness group diverges into two independent high fitness lineages (H1 and H2), each characterized by distinct CN profiles. Notably, the second principal component values of H1 and H2 are similar, suggesting that shared aneuploidies may underlie the observed fitness jumps despite their overall divergent evolutionary path in the CN space.

### Differential aneuploidy landscapes distinguish high- and low-fitness WGD pop-ulations

As certain chromosomal copy number alterations have been reported to confer fitness effects in cancer^50^, we investigated whether LeafRank’s predictions could facilitate integrative genomic analysis to identify specific SCNAs underlying the observed fitness jumps. We reason that if a specific SCNA enhanced cellular fitness, it would be enriched, or recurrently observed within high fitness groups as compared to their low fitness counterparts.

To this end, we identified SCNAs associated with the fitness differences between the cells predicted with high fitness and those with low fitness across the ten truncal WGD HGSOC tu-mors (see Methods). In metastatic tumor OV-087 (omentum), for example, the fitness grouping closely correspond to the SCNA lineage structure revealed by the phylogenetic tree (Figure 6). The highest-fitness group exhibits specific SCNAs relative to the rest of sampled cells, including arm-level aneuplodies such as loss of Chromosome 10q (Chr10q) and gain of entire Chr18, as well as segmental gain in Chr9p and partial loss of Chr8q. Notably, Chr10q shows the strongest significance, harboring the canonical tumor suppressor gene *PTEN*.

**Figure 6.**
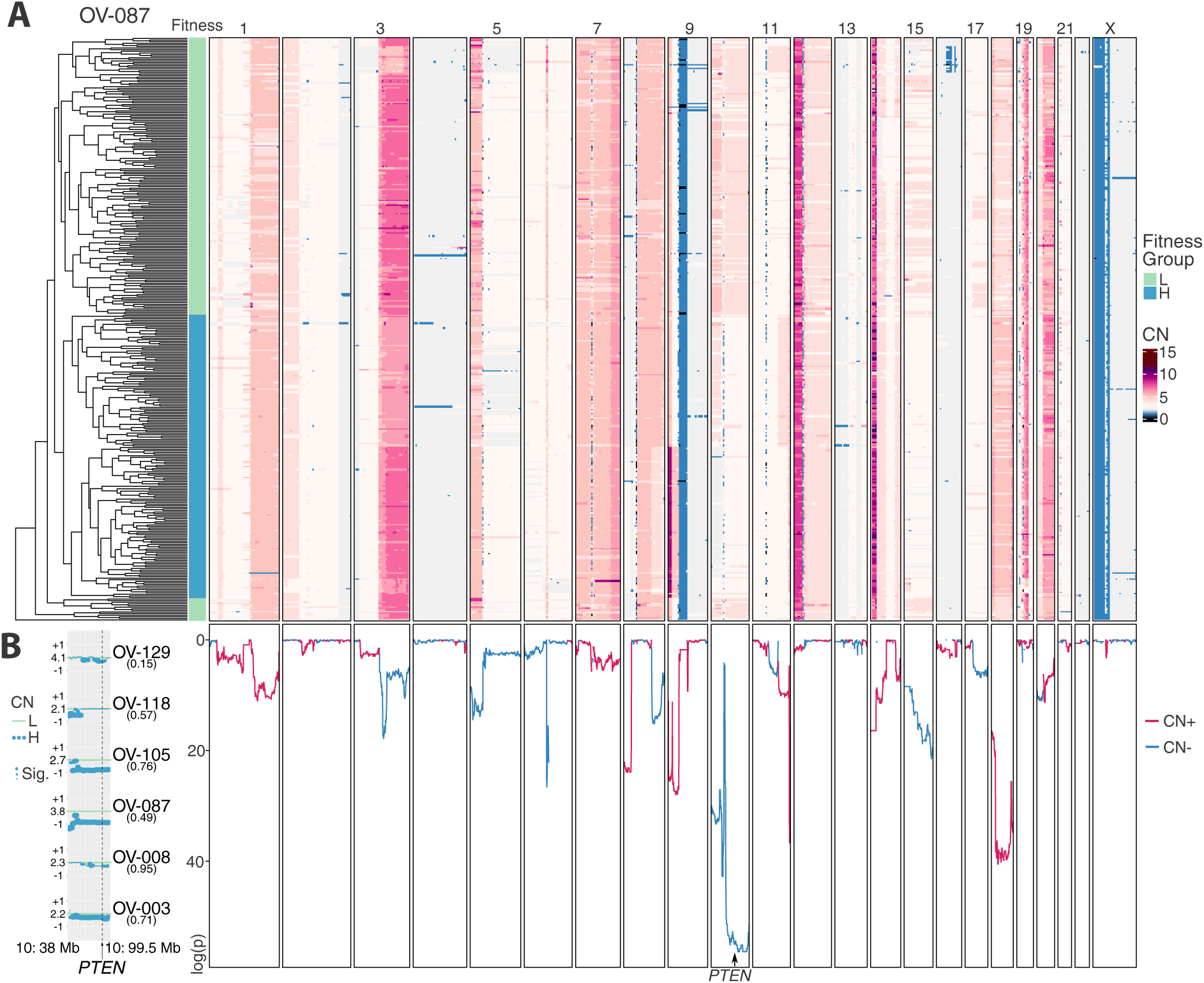
Detecting SCNAs associated with fitness differences. **(A)** In OV-087, copy number pro-files of sampled cells (right, color-coded heatmap) are shown alongside an ultrametric tree (left) rescaled based on CN events. A dichotomous grouping of single cells according to inferred fitness is displayed between the tree and the CN profiles. **(B)** The panel below the CN heatmap shows the log-transformed p-values from Wilcoxon tests assessing each genomic bin for differences in copy number states be-tween high-fitness (H) and low-fitness (L) cells. The left panel highlights a recurrent CN region on Chr10 (p11.1–q24.2) that exhibits a significant and consistent CN reduction in high-fitness cells across six HG-SOC tumors, harboring the *PTEN* locus. Details regarding the identification of this region are provided in the Methods. For each patient, the average CN of genomic bins within this region for low-fitness cells is plotted as a green baseline. Blue dots represent the CN difference between the high- and low-fitness cell groups for each individual genomic bin. The fraction of high-fitness cells is indicated in parenthe-ses. Additional recurrent, fitness-associated CN regions are presented in the Supplemental Text and Figure S5B.

Most fitness associated SCNAs are patient specific, rarely recurring in more than three patients. Moreover, we observed that while a specific chromosomal region may present fitness associated gains in one patient, it can show fitness-associated losses in another (Figure S5). This suggests that the fitness effect of SCNAs are highly context-dependent and influenced by patient-specific fitness landscapes^51^. However, reduced CN for Chr10 (p11.1 - q24.2), which contains *PTEN*, was recurrently associated with fitness increases (p < 0.05) in six patients (Figure 6). This recurrence suggests the consistent fitness effect conferred by SCNAs affecting this region during post-WGD evolution in HGSOC, distinguishing it from the more context dependent aneuploidies.

We also identified additional chromosomal regions exhibiting recurrent fitness associated SCNAs across patients (Figure S5). In particular, we identified gains involving Chr12 (p13.31 - p11.23), which contains *KRAS*, consistent with recurrent amplifications reported in large cohort studies^52–54^. Moreover, fitness associated losses on Chr6 (q15 - q16.3) have been previously linked to chemoresistance^54^, suggesting that this alteration may contribute to preexisting treatment resistance by conferring a selective advantage in HGSOC. These findings suggest that the heterogeneous SCNA landscape in HGSOC often obscures consistent CNA signals. The high-resolution relative fitness inference provided by LeafRank therefore offers a powerful framework for uncovering these aneuploidy-driven fitness pattern that might otherwise remain hidden in standard genomic analysis.

### Fitness ranking reveals equally competitive WGD**^−^** lineages in Subclonal WGD Tumors

To investigate the fitness effect of WGD itself, we analyzed three subclonal-WGD HGSOC patient tumors where WGD^+^ and WGD^−^ cells coexist. In these phylogenetic trees, constructed from SCNA events by MEDICC2^28^, WGD^+^ lineages consistently exhibit longer branch lengths than WGD^−^ populations, mirroring the pattern observed in our simulated WGD tumors. This observation is consistent with prior findings that WGD increases the SCNA rate^39–41^. To account for this effect, we recalibrated branch lengths using our modified *chronos* algorithm (Methods) to generate WGD-adjusted ultrametric trees (Figure 7A). Because the ultrametric reconstruction may depend on assumptions regarding WGD branch classification and rate modeling, we first assessed how well the proposed WGD-state dependent molecular clock model fit to the observed branch length. Additionally, we examined the robustness of downstream inferences under alternative calibrations schemes (Supplemental Text).

**Figure 7.**
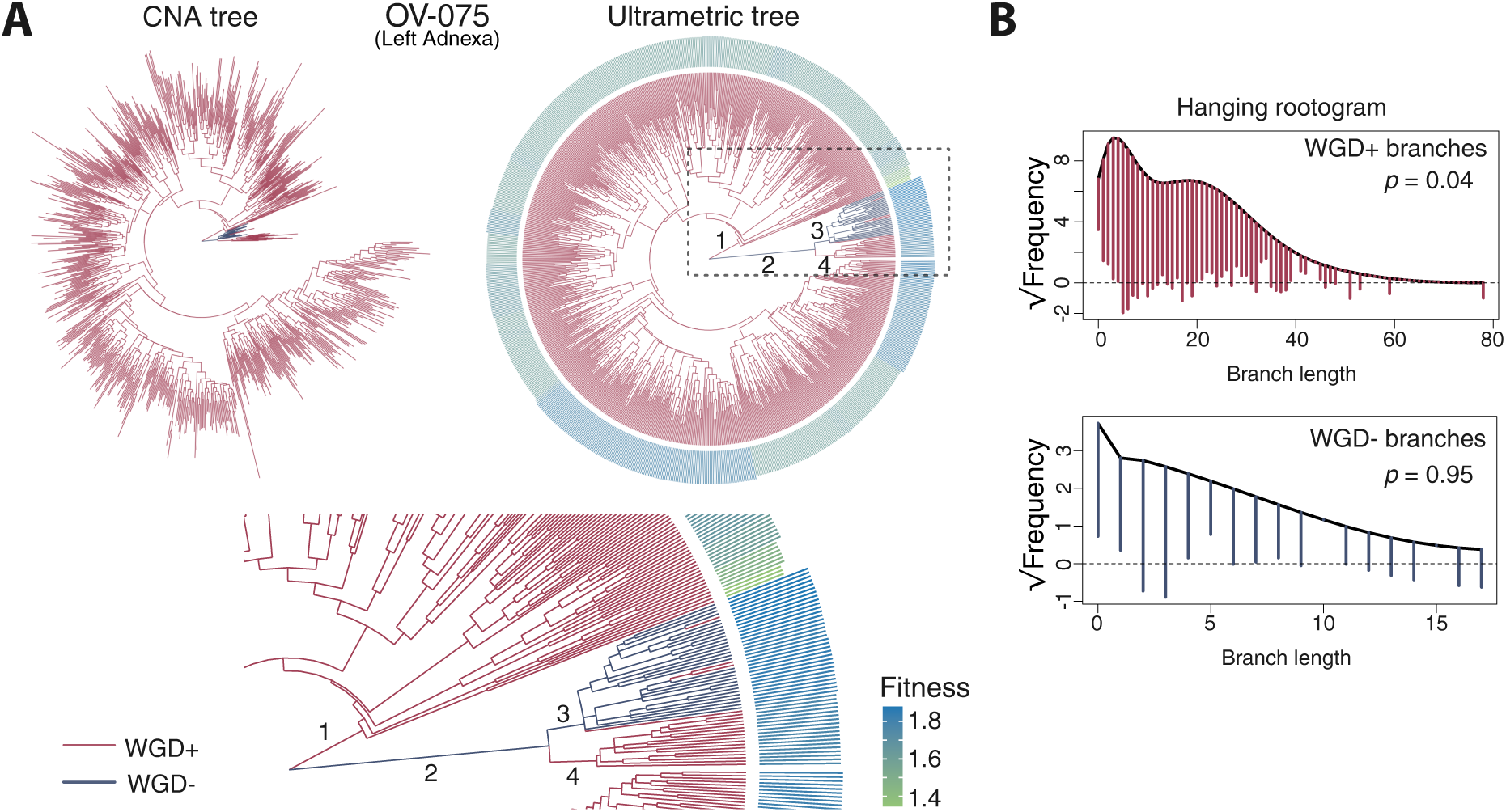
Subclonal WGD cells do not consistently exhibit higher fitness than their diploid counterparts. **(A)** The CN-event phylogenetic tree (left) derived from MEDICC2 and the rescaled ultrametric tree (right) for patient tumor OV-075 are shown, which include both subclonal WGD^+^ and WGD^−^ cells. The lower panel provides a zoomed in view of the ultrametric tree, highlighting the contrast in inferred fitness between WGD^+^ and WGD^−^ cells. **(B)** Hanging rootogram illustrating the performance of ultra-metric tree conversion for both WGD^+^ and WGD^−^ branches. The black solid line denotes the expected frequency distribution of branch lengths calculated from the rescaled ultrametric tree using inferred pa-rameters under a Poisson model (See the Methods and Supplemental Text), whereas the color-coded hanging bars represent the observed frequency distribution from the MEDICC2 tree. Frequencies are square-root scaled.

To assess whether the Poisson-based branch length model underlying our modified *chronos* algorithm remains applicable to patient tumors, we compared the expected branch length distri-bution under the Poisson assumption, derived from the timing information in the WGD-adjusted ultrametric tree, with the observed branch lengths from the original MEDICC2 tree (see Sup-plemental Text). Although SCNA events are not strictly independent, as each event alters the available DNA substrate for subsequent alterations, the Poisson model provides a good over-all approximation of the observed branch length distribution (Figure 7B). Differences between expected and observed frequencies are generally small and centered around zero across most branch lengths, with deviations primarily observed among shorter branch lengths (< 15 SCNA events). Specifically, the model overestimates the frequency of branches with no or few SCNA events, whereas the MEDICC2 tree shows an enrichment of intermediate branch lengths corresponding to approximately five events. This discrepancy may reflect punctuated SCNA acquisition in WGD^+^ lineage, as previously proposed in WGD tumors^55^, or limitations in the sensitivity and accuracy of SCNA detection from scDNA-seq data^30^.

Using the WGD-adjusted ultrametric trees, LeafRank revealed that WGD^+^ cells do not exhibit a uniform fitness advantage over their WGD^−^ counterparts at the time of sampling. This finding was consistent across multiple subclonal WGD tumors and remained robust to alternative calibration assumptions for branch classification and rate modeling (see Supplemental Text and Figure S7). In primary tumor OV-075 (left adnexa), two lineages diverged from the founder: one giving rise to WGD^+^ descendants (branch1, Figure 7), and the other initially remaining WGD^−^ (branch2) before acquiring a subsequent WGD event in a sublineage (branch4). Notably, although branch2 is shorter than branch1 in the original MEDICC2 tree, it becomes substantially longer in the WGD-adjusted ultrametric tree. This reflects the lower SCNA rates in the WGD^−^ lineage, which requires more elapsed time to accumulate a comparable number of SCNAs events. Correspondingly, the terminal branches descending from branch2 are relatively short, indicating that this lineage expanded to a detectable population size over a shorter evolutionary time interval. This “late but rapid” branching results in a higher predicted fitness. Within this lineage, the newly emerged WGD^+^ subclone (branch4) constitutes a smaller population yet displays a branch length comparable to its WGD^−^ sibling lineage (branch3), suggesting no relative fitness advantage associated with WGD at this bifurcation. In contrast, the WGD^+^ lineage derived from branch1 occupies a large fraction of the tumor sample but displays substantial internal fitness heterogeneity, characterized by both low-fitness populations arising at earlier stages and higher-fitness populations that emerged from the continuously expanding WGD^+^ lineage. This is consistent with observations in the truncal WGD tumors.

These results demonstrate that while WGD is a landmark recurrent event in HGSOC, its occurrence does not necessarily confer a strong fitness benefit. Instead, its prevalence in HG-SOC can be explained by enhanced long-term evolvability, such as an increased rate of driver aberrations or a greater robustness to microenvironmental perturbations. These findings are consistent with prior studies in cancer^56^ and microorganisms^57^. Importantly, LeafRank allows for the direct assessment of fitness of this event in patient tumors, highlighting its potential to address unresolved questions in cancer genomics.

## DISCUSSION

Cancer cells can exhibit substantial intratumor heterogeneity in fitness, and single-cell phylogenetic trees reconstructed from SSNVs and SCNAs can encode structural patterns that reflect this diversity. In this study, we introduce **LeafRank**, a probabilistic framework for ranking cellular fitness from single-cell phylogenetic trees. Unlike existing fitness-estimation approaches in cancer^32,33,38^, which often focus on the absolute fitness effects of specific genomic alterations or predefined subclones, LeafRank estimates the relative fitness ranking of sampled cells at the leaves of the tree. By combining a multitype branching-process model with a message-passing algorithm, LeafRank integrates information encoded in the single-cell phylogeny, including branch lengths and bifurcation patterns, to produce robust cell-level fitness estimates. Our results show that, during cancer clonal expansion with discrete levels of selective advantage, tree structure contains critical information about the relative fitness of sampled cells. This fitness signal is captured by the branch lengths and bifurcation patterns of the phylogeny. Importantly, we found that the resulting fitness rankings are largely robust to the specific choice of input parameters in the underlying branching-process model. This observation is consistent with previous findings in asexual populations of constant size with clonal interference^19,42^, where reliable fitness rankings were obtained across a broad range of parameter values. However, phylogenetic patterns reflecting selection in expanding cancer populations may differ substantially from those in fixed-size populations. Despite these differences, the observed consistency suggests that tree structure can encode robust relative-fitness information across distinct evolutionary regimes, enabling fitness inference even when precise parameter values are not fully characterized.

LeafRank provides a generalizable framework for fitness inference from single-cell phylogenies. First, it does not assume uniform fitness within a subclone, allowing selection within a given lineage to be resolved. Second, the message-passing algorithm provides an efficient framework for propagating information along the tree and can flexibly incorporate prior knowledge or model assumptions. Third, because LeafRank provides fitness estimates for every sampled cell, it enables high-resolution comparisons of populations grouped not only by genomic alterations but also by other functional features, such as gene-expression profiles or epigenetic states, when multiomic data are available. This flexibility facilitates the study of the determinants of growth-rate variation in cancer. For example, by overlaying transcriptional signatures onto fitness rankings, one can identify pathways associated with the clonal expansion of high-fitness lineages, thereby linking genomic architecture to cancer phenotype.

A key challenge when inferring fitness in clinical samples is the reliance on an evolutionary molecular clock. In patient tumors with CIN, lineages can exhibit accelerated rates of SCNAs and SSNVs, severely violating linear temporal assumptions^39–41^. To resolve this, LeafRank incorporates a tree-rescaling strategy capable of navigating the piecewise linear tempos introduced by macro-evolutionary events. By decoupling the aberration rates of WGD^+^ and WGD^−^lineages and optimizing a joint likelihood function, the framework effectively charts independent rate heterogeneity onto a common temporal scale. This normalization allows LeafRank to robustly analyze phylogenies that would otherwise confound standard molecular-clock models.

By analyzing recently published single-cell WGS data from HGSOC patients^44^, LeafRank identified cell populations with distinct selective advantages within tumors characterized by truncal WGD. While these tumors were predominantly sampled from metastatic sites, signs of selection were also evident in a primary tumor (OV-118) as well as in the subclonal WGD+ lineages of two additional primary cases. This indicates that the inferred selective advantages may reflect ongoing evolutionary pressures operating both early in tumor development and post-dissemination, potentially facilitated by WGD-associated CIN. This detectable selection underlying HGSOC expansion contrasts with previous bulk-sequencing studies of colorectal cancer that proposed an ‘effective-neutral’ model for some patients^12,58^. This divergence likely highlights the enhanced resolution of scDNA-seq, which unmasks subclonal fitness differences that are otherwise obscured by bulk averaging. Furthermore, while early bulk studies focused primarily on SSNVs in diploid regions, our framework leverages SCNA-based phylogenies to capture growth dynamics. We note, however, that when bulk sequencing is extended to multi-region sampling, subclonal selection has indeed been detected in colorectal cancers with CIN^13^. The high-resolution fitness mapping provided by LeafRank enabled the identification of specific post-WGD SCNAs associated with increased inferred fitness. Notably, five patients exhibited recurrent Chr10q loss encompassing the tumor suppressor *PTEN*. This suggests that *PTEN* haploinsufficiency may drive subclonal expansion after WGD in HGSOC. Our observation is consistent with the “dosage-sensitivity” model of tumor suppressors, in which partial reduction of *PTEN* dosage through large-scale chromosomal deletion may disrupt the dosage balance of the PI3K/AKT signaling pathway, potentially conferring a selective advantage. Beyond Chr10q loss, several other SCNAs were recurrently associated with high-fitness subclones, including Chr12p amplification and Chr6q deletion. Chr12p was reported to be recurrently amplified in ovarian cancer^52,53^ and Chr6q loss has been linked to chemo-resistance^54^, highlighting the biological relevance and clinical utility of the fitness rankings inferred by LeafRank.

Although WGD may buffer against deleterious aberrations^59^, our results suggest that its prevalence in HGSOC cannot be explained by an immediate fitness advantage. In tumors containing both WGD^+^ and WGD^−^ lineages, WGD^+^ cells do not universally outcompete diploid cells. These results were derived from rescaled trees under the assumption that WGD status is maintained from the start of the branch containing the inferred WGD event. We note that the precise temporal placement of WGD within this branch remains challenging. However, even when we conservatively assign the initial branch as WGD^−^, effectively modeling WGD as arising at the subsequent bifurcation, the inferred fitness of early WGD^+^ cells consistently remained lower than that of WGD^−^ lineages (Figure S7). Taken together, LeafRank’s predictions in HGSOC tumors are consistent with existing hypotheses and empirical findings in cancer and other biological contexts^56,57^, suggesting that WGD primarily confers long-term evolutionary advantages rather than immediate fitness gains.

In summary, by integrating information encoded in single-cell phylogenies, LeafRank captures fine-scale evolutionary dynamics during cancer growth. It provides a generalizable tool for translating tree structure into relative fitness rankings of single cells, offering a powerful framework for identifying the aberrations that drive cancer clonal expansion.

## Limitations

LeafRank has several limitations. First, it is most informative in evolutionary regimes in which subclonal selection drives heterogeneous fitness levels across sampled cells. In such settings, the tree structure deviates from expectations under neutral evolution and contains a detectable signal of differential growth. By contrast, LeafRank is less informative for tumors evolving under strict neutrality, or for tumors that appear effectively neutral after a recent selective sweep, because sampled cells may show little residual tree-structural evidence of fitness differences. In these cases, LeafRank may return nearly uniform or weakly resolved fitness rankings rather than identifying distinct high-fitness lineages^12^.

Despite the utility of our tree rescaling framework, LeafRank’s performance remains bound to the quality of the input ultrametric tree. While using accumulated somatic aberrations serves as a pragmatic molecular-clock proxy, the underlying mutational processes can fluctuate wildly across lineages and over time. Consequently, in tumors with intricate subclonal WGD histories, our modified *chronos* implementation may only partially mitigate severe rate distortions, potentially reducing the reliability of downstream fitness rankings. Resolving these topological ambiguities in highly rearranged genomes remains an open challenge. Ultimately, overcoming this limitation will likely depend on integrating emerging multi-omic profiling^60^ or CpG methylation-based lineage-tracing methods^61^, which promise to decouple physical time from mutational density and yield more robust input phylogenies.

Lastly, LeafRank assumes a tumor population that is still expanding in size at the time of sampling. Our spatial simulations indicate that LeafRank tolerates spatial structure and local crowding during this expansion phase, recovering accurate rankings even when the perfectly mixed assumption is violated (Figure 4G). However, once net population growth ceases, because the tumor has approached carrying capacity, stabilized, or entered a prolonged regime of population stasis, the genealogy increasingly resembles that of a constant-size population and loses the expansion signal LeafRank relies on, reducing the informativeness of the inferred rankings. Capturing this saturated regime will require alternative generative assumptions, such as logistic or otherwise density-dependent branching processes.

## METHODS

### LeafRank: model setup

A binary phylogenetic tree, denoted by F, records the evolutionary history from a common ancestor cell to all N–1 internal nodes and N leaf nodes. Such a tree can be reconstructed using algorithms such as neighbor-joining^62^ or MEDICC2^28^ based on single-cell genomic profiles. As illustrated in Figure 1, we denote the tree topology as T, which includes the branch topology (the topological information describes the sibling relationships among cells) and the branch length information (the evolutionary distance, measured by the number of SSNVs or SCNAs, between ancestor and descendant cells). Note that under an assumed genomic aberration rate, the branch length information can be converted into estimates of the evolutionary time elapsed along each branch.

Temporal calibration of branch lengths often relies on the molecular clock assumption^63^, under which genomic aberrations are assumed to accumulate at a relatively constant rate over time. Under this assumption, we denote the constant neutral aberration rate by *µ*. While this assumption is widely adopted in the context of SSNVs, it may not necessarily hold for SC-NAs. For example, SSNV and SCNA rates can increase substantially following WGD^39–41^. In a later subsection and in the Supplemental Text, we further discuss our approach to this issue, particularly with respect to WGD events, and provide corresponding validation analyses.

To model the underlying cellular evolutionary dynamics, we employ a multitype branching process model^45^. Under this model, we assume that there are V cell types, each characterized by distinct birth rates (b_i_) and death rates (d_i_), where i ∈ {1, · · ·, V}. The cell types are ordered by increasing fitness such that b_i+1_ > b_i_. For simplicity, we assume the death rate d_i_ is constant across all cell types. We use a backward-time convention for the phylogenetic tree, with the sampled cells observed at time t = 0 and the root located at time t = T. Thus, T denotes the elapsed time between the common ancestor and the sampled population. The biological process starts from a single type-1 cell at the root and evolves toward the sampled population. To model driver events, we further assume that a type i cell can divide into one type i and one fitness-advantaged type i + 1 cell at rate *ν*, which we refer to as the driver aberration rate. Notably, this rate *ν* is assumed to be much smaller than the neutral aberration rates *µ*, reflecting the rarity of driver events in cancer evolution. At the sampling time, we assume a uniform sampling scheme in which each living cell has an equal chance *ρ* ≪ 1 of being sampled. In practice, this value can be approximated as the ratio of sampled cell counts to the total population size when the population is sufficiently large. In the following *in silico* study, we sample 250 cells from a population of one million cells, resulting in an approximate sampling probability of 0.00025.

To model the cellular fitness states across the tree, we denote the identities of the leaf and internal nodes as random variables **v**_X_ = {v_x1_, · · ·, v_xN_ } and **v**_Y_ = {v_y1_, · · ·, v_yN–1_ } respectively, each taking values in the state space {1, · · ·, V}. These quantities are not directly observable.

Instead, LeafRank is designed to infer the probability distribution of v_x_ for each leaf given the tree structure and the model configuration. Specifically, we compute the probability density of the phylogenetic tree F:

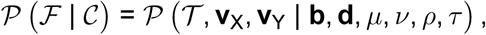

where **b**, **d** are vectors for birth and death rate for each type, respectively, and *τ* denotes the scaling factor for time elapse.

### Computing marginal fitness distribution of sampled cells

The goal of LeafRank is to rank sampled cells at the leaves according to their distributions over fitness types, conditioned on the observed phylogenetic tree topology T and model configuration C. For a sampled cell x ∈ X, this marginal distribution is

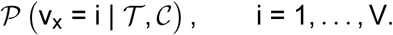

Equivalently, it can be obtained from the joint distribution over all leaf and internal-node type assignments by summing over all latent configurations consistent with v_x_ = i:

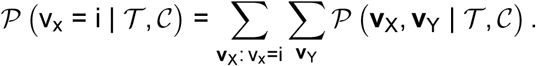

Direct evaluation of this expression is computationally inefficient, because the number of possible assignments of latent fitness types increases exponentially with both the number of fitness states and the size of the tree.

To compute the marginal fitness distribution of each sampled cell efficiently, we use *message passing*, also known as *belief propagation* in statistical physics and information theory^64^. Message-passing methods have previously been applied in evolutionary settings, including analyses of seasonal influenza A/H3N2^42^. In our setting, however, the application is not immediate: the local quantities required for message passing are not available as simple discrete factors on the tree, but instead must be derived from the continuous-time multitype branching process that defines the phylogenetic density.

Specifically, we exploit the tree structure and express the distribution of the type at node n as the product of two complementary terms: an upward message from the descendant subtree T_n–_ and a downward message from the remainder of the tree T_n+_,

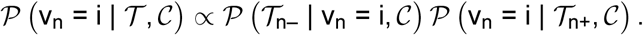

This factorization avoids explicit enumeration of all latent fitness-type assignments on the phylogeny. The upward pass summarizes information contributed by descendants of node n, whereas the downward pass propagates information from the root together with information from the sibling side of the tree (Figure S1 and Figure 1).

Importantly, this message-passing formulation cannot be applied directly in our model. We therefore first decompose the phylogenetic density into two elementary building blocks: branch propagators, which transport type-dependent densities along individual branches, and bifurcation propagators, which combine the two daughter subtrees at each internal node. This decomposition is the key step that converts the global phylogenetic density into a collection of local recursive operations that can be assembled by message passing.

For the branch propagator, we adopt the BiSSE model^65^ to compute the type-dependent densities along each branch. Specifically, we track a probability D_n,v_(t) defined as the probability that a lineage starting at time t in state v evolves into the sampled clade subtended by node n. For each state i ∈ {1, · · ·, V}, the quantity D_n,i_(t) is propagated backward in time along the branch, from the time of the immediate descendant node t = t_d_ to that of the ancestor node t = t_a_. Within an infinitesimal time interval, at most one cellular event occurs, such as a division event in which a type i cell produces two type i cells, denoted by i → {i, i}. Under this framework, D_n,v_(t) can be computed by solving:

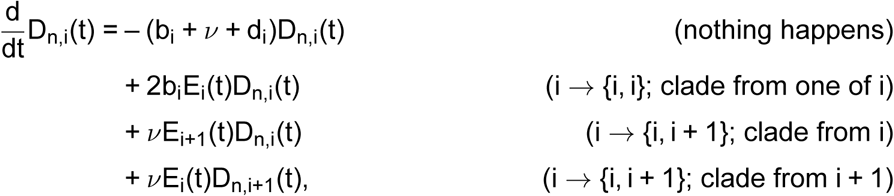

where E_i_(t) is defined as the probability that an individual type i cell has no sampled descendants after time t. Note that when i = V, driver aberration terms will disappear. A crucial observation is that, these differential equations governing the branch densities form a time-varying linear system. This linearity allows each branch to be solved independently under basis initial conditions, and the solution for arbitrary inputs can then be reconstructed by superposition. As a result, the overall computation separates into independent branch-level and bifurcation-level components that can be evaluated efficiently and then combined through the upward and downward recursions. In this way, we obtain the marginal fitness distribution at every leaf without enumerating all internal-node state configurations. For a detailed derivation and formal definition of the bifurcation propagator, we refer the reader to the Supplemental Text.

### Benchmarking LeafRank using simulated data

To validate LeafRank through *in silico* experiments, we simulated both non-spatial and spatial virtual tumors^43^. In this subsection, we document the detailed simulation settings and the corresponding inference configurations for LeafRank, organized according to each experiment conducted.

### Non-spatial simulation

In the non-spatial simulation, cellular evolutionary dynamics are modeled using a multi-type branching process. Stochastic branching events are simulated using the classical Gillespie algorithm^66^, assuming exponentially distributed waiting times determined by the corresponding event rates. Each rate represents the instantaneous probability per unit time that a given event occurs. Specifically, we consider five classes of stochastic events: cell birth (b_i_), cell death (d), driver aberration (*ν*), passenger aberration (*µ*), and WGD (*ω*). For simplicity, we assume the existence of eight discrete fitness states. These states share a common death rate d = 0.18 but differ in their respective birth rates, defined as b_i_ = 0.2 × 1.2^i–1^. This parameterization models birth rates that scale multiplicatively by a factor of 1.2 across successive phenotypic levels. Driver aberrations occur at rate *ν* = 0.0001 and induce irreversible transitions from lower- to higher-fitness types. To construct DM-based phylogenetic trees, we additionally incorporate passenger aberrations at a substantially higher rate *µ* = 0.18 to record pairwise cellular divergence. Upon acquisition of a WGD event at rate *ω* = 0.0001, both parameters *ν* and *µ* are doubled. The resulting distance matrix is subsequently used as input for phylogenetic reconstruction via neighbor joining^62^.

Using these parameter settings, we simulated 50 diploid tumors (*ω* = 0) and 50 WGD-aware tumors (*ω* = 0.0001). Each simulation was terminated when the number of viable tumor cells reached one million, after which point 250 cells were randomly sampled from the surviving population for downstream analyses. Due to the stochastic nature of the branching process, tumor extinction can occur during simulation. In such cases, the simulation was restarted from a single type-1 cell. All simulations and sampling procedures were implemented in MATLAB using the *PriorityQueue* data structure^67^.

### Spatial simulation

We extend our existing single-cell-based spatial tumor growth model, **Comet**^43^, to simulate the virtual tumors with ongoing WGD. In the spatial model, tumor expansion starts with a single founder cell located in the center of a three dimensional lattice with the Moore neighborhood. Cells divide at rate b and die at rate d per unit time. The waiting time between two consecutive division events is assumed to be exponentially distributed with mean 1/b. We set the b for the founder cell at 0.25 by assuming that the average cell cycle is four days. The death rate is chosen at 0.2475 to achieve a death-to-birth ratio at 0.99, mimicking the slow growth of primary tumor. When a cell divides, one of its daughter cells stays at the original position on the grid, whereas the other fills one of the neighboring empty sites uniformly at random. When a cell dies, it is removed from the lattice. To consider spatial constraints in solid tumors, the actual birth rate of a cell is adjusted by the proportion of empty neighboring sites.

To model genome evolution, **Comet** assigns aberrations to abstract genomic coordinates. Passenger aberrations arise at rate *µ* = 0.15 per cell division, while driver aberrations arise at rate *ν* = 0.0001. WGD occurs at a rate 0.0005. Upon WGD, both *µ* and *ν* are doubled. To allow aberrations, particularly SCNAs, to overlap, the coordinate space is restricted to a finite pool: 5 × 10^7^ for passengers and 2 × 10^5^ for drivers. As in the non-spatial model, each driver confers a baseline selection coefficient of 0.2, but with additional variability introduced by Gaussian noise with a standard deviation of 0.1. For this study, once the population reached one million cells, we randomly sampled 250 cells from the virtual tumor. The TT was obtained by tracing the specific lineage and birth time of each cell. Phylogenetic trees were then reconstructed using the aberration profiles from these sampled cells.

### Ranking stability under subsampling

To evaluate the relationship between LeafRank’s ranking stability and its estimation accuracy, we performed systematic downsampling experiments (Figure 2D and Figure 4H). For each virtual tumor, we randomly subsampled 80% of the original sampled dataset 50 times. We then inferred relative fitness values for each independent subsample and computed the correlation between the resulting rankings and the corresponding ground-truth rankings. Inference accuracy was evaluated by computing the mean correlation coefficient of the subsampled rankings against the ground truth. Concurrently, ranking instability was quantified by calculating the variance of the correlation coefficients between the subsampled rankings and the baseline ranking obtained from the complete, non-subsampled population.

### Ovarian cancer patient data overview

The single-cell high-grade serous ovarian cancer (HGSOC) patient data reported in McPherson et al.^44^ contains 70 HGSOC samples derived from 41 patients. The samples are collected pre-treatment from primary (left or right adnexa) or metastatic sites (e.g., omentum, peritoneum, bowell). The scWGS are generated using direct library preparation (DLP+) platform^26^, which constructs single cell sequencing libraries without pre-amplification. By improving coverage uniformity and minimizing amplification bias, this platform enables reliable detection of SCNAs at single-cell resolution. To reconstruct WGD-aware phylogenetic trees, the processed haplotype-specific single-cell CN profiles were previously analyzed using MEDICC2^28^. MEDICC2 infers evolutionary trees based on a minimum-event distance metric that accounts for genomic contiguity by allowing overlapping CN event rather than treating adjacent loci as independent. For the present study, we obtained and analyzed these pre-computed MEDICC2 trees directly from the original publication^44^. The R package *ape* (Version 5.8.1) is used to store and analyze the phylogenetic tree.

Among 41 patients, the authors categorized the patient samples into 4 groups: 1. Truncal WGD (21 patients), 2. Parallel WGD (2 patients), 3. Subclonal WGD (5 patients), 4. Un-expanded WGD (11 patients). We mainly focused on a subset of ten patients classified as Truncal WGD (Figure S3 and Figure S4) and three patients as Subclonal WGD (Figure 7 and Figure S6), where cells were sampled from the same anatomical site. Tumors exhibiting Truncal WGD display leaf-to-root branch lengths that remain relatively uniform across sampled cells, thereby satisfying the baseline assumption of a constant SCNA accumulation rate. In contrast, for tumors showing Subclonal WGD, the WGD^+^ subpopulations typically present more protracted developmental histories, which are explicitly modeled and accounted for using a Poisson framework (see next section). Cases of Parallel WGD were excluded from further analysis as none met the identical anatomical site requirement. Cases of Unexpanded WGD were excluded because the WGD^+^ cells has not reach sufficient population size.

### Using LeafRank to analyze distance-matrix based phylogenetic tree

Our proposed algorithm, LeafRank, assumes a reconstructed phylogenetic tree whose branch lengths reflect the elapsed time between ancestral and descendant nodes. For samples collected at a single time point, an ultrametric tree therefore provides the ideal input. In practice, however, phylogenetic trees are more commonly reconstructed using DM-based methods, such as neighbor-joining or MEDICC2^28^, because aberration distances can be directly inferred from genomic data. Under a traditional molecular clock assumption, where aberrations accumulate at a constant rate, genomic divergence is directly proportional to elapsed time. This alignment allows distance-based trees to be interpreted within a temporal framework and utilized directly by LeafRank. However, this assumption is frequently violated by macro-evolutionary events such as WGD, which can induce CIN and trigger heterogeneous aberration rates across distinct lineages. To address this limitation, we developed a procedure to recover ultrametric trees under a state-dependent molecular clock model, accompanied by goodness-of-fit diagnostics and a parameter configuration framework for the transformed trees.

### Converting a DM tree into an ultrametric tree

To estimate an ultrametric tree, we adopt the *chronos* function^46^ in R. The *chronos* function constructs an ultrametric phylogenetic tree from DM tree under a molecular clock assumption coupled with a Poisson model. Specifically, it assumes the number of aberrations follow the Poisson distribution with mean proportional to the product of evolutionary time and a constant aberration rate. Using maximum likelihood scheme, it jointly infers a constant aberration rate and branch-specific evolutionary time to maximize the likelihood of observing aberration distances along each branch (see Supplemental Text).

To capture the potential heterogeneity in aberration rates between WGD^+^ and WGD ^−^ populations, we propose a state-dependent molecular clock model. In this model, we classify tree branches as WGD^−^ and WGD^+^ branches according to the WGD status of its descendant node. In other words, we make the following assumption:

**Assumption** (*Anc.*). For each WGD^+^ *clade, the WGD event is assigned to the ancestral end of the branch leading to the earliest node inferred to be WGD*^+^*. Consequently, the branch leading to this earliest WGD*^+^ *node is labeled as WGD*^+^*, together with all branches descending from this node*.

For an internal node, if both daughter lineages are WGD^+^, then the parent node is inferred to be WGD^+^, under the assumption that WGD status is irreversible. This irreversibility assumption is consistent with the minimum-event framework used by MEDICC2^28^: a return from a genome-doubled state to a diploid-like state would require many copy-number losses and is therefore not favored under a parsimonious copy-number-event reconstruction.

We then modified the ultrametric tree-inference framework using the following state-dependent Poisson model:

**Assumption** (*Poi.*). The WGD^+^ *population accumulates SCNAs according to the same Poisson process as the WGD*^−^ *population, but with an approximately twofold higher aberration rate*.

This assumption does not impose an exact twofold relationship between the alteration rates of WGD^+^ and WGD^−^ branches. Instead, the two rates are estimated from the data.

Under these assumptions, we formulated and solved a modified maximum likelihood estimation problem (see Supplemental Text) to jointly estimate two aberration rates and the corresponding adjusted branch lengths of the ultrametric tree. The optimization problem was solved in MATLAB (Version R2025b) using Optimization Toolbox function *fmincon*^68^.

### Using rootograms to evaluate the reconstructed ultrametric tree

To evaluate the goodness of fit of the rescaled ultrametric tree, we employed rootograms^69^, which visually expose discrepancies between observed and expected count distributions. Specifically, we utilized hanging rootograms to compare the empirical distribution of DM-tree branch-lengths against the theoretical distribution expected under the fitted Poisson model. In a hanging rootogram, the bars are suspended from the fitted frequencies, such that deviations from the zero-reference line highlight systematic discrepancies. Beyond visual inspection, we examined the sign pattern of the residuals across branch-length intervals to assess whether unexplained structural dependencies remained. This analysis was performed using a runs test, implemented with *runs.test*. Under a well-fitting model, positive and negative residuals are expected to alternate without systematic structure, whereas extended runs of residuals with the same sign may indicate model misspecification. We note that this diagnostic relies solely on residual signs rather than magnitudes. It may also be affected by dependence among branch-level observations. Therefore, the runs test should be interpreted as an informative heuristic of model adequacy rather than a definitive, formal goodness-of-fit test.

### Preset input parameter configuration

Once an ultrametric tree is reconstructed, we provide a set of input parameter configurations, accompanied by justifications derived from the underlying branching-process model. As detailed in the Results section, these model parameters are highly interdependent and jointly determine the informativeness of the inferred fitness distribution. Rather than prescribing a universal parameter configuration, we present a principled framework for selecting individual parameters while conditioning on the remaining ones. We illustrate this procedure using the time-scale parameter *τ* .

Consider a ultrametric tree whose root-to-leaf distance is scaled to 1. Under the convention that the internal model time is obtained by dividing normalized branch lengths by *τ*, the time-scale parameter can be interpreted as 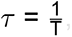, where T denotes the expected elapsed time from the MRCA to the sampled leaves. We estimate T under a given parameter configuration as follows. Suppose the tree contains m sampled leaves and the sampling probability *ρ* is known. The total population size can then be approximated by 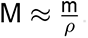.

Given the birth rates b_i_, death rates d_i_, and driver-event rate *ν*, we construct the mean matrix **A** for the multitype branching process. Under the driver-birth model used by LeafRank, and using a row-vector convention, this matrix has entries

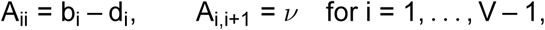

with all other entries equal to zero. Under this model, the expected total population size at elapsed time T satisfies

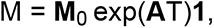

where **M**_0_ is the initial population vector and **1** is the all-ones column vector, which sums over all fitness phenotypes at time T. Assuming that the MRCA consists of a single type-1 cell, the initial condition is

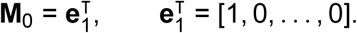

Solving the equation above yields the estimate of the expected elapsed time T from the MRCA to the sampled leaves, and therefore determines the corresponding time-scale parameter *τ* .

A similar strategy can be used to estimate other parameters when the time scale is fixed. In the analysis of HGSOC patient data, we adopted this approach to determine the time-scale pa-rameter under a fixed sampling probability and a specified fitness-phenotype configuration. As shown in Figure S3 and Figure S4, the resulting parameter configurations produced informative inferred fitness distributions across all analyzed tumors.

### LeafRank reveals fitness-advantageous aneuploidy pattern

#### Genomic coordinates and chromosome binning scheme

To identify the genomic coordinates of *PTEN* and other target genes, we utilize the *hg19* refer-ence genome build. The genome is segmented across 23 chromosomes into 6,087 contiguous bins, with a uniform bin size of 500 kb. Consequently, both the continuous copy number read-depth matrix and the discrete copy number state table possess dimensions of N_bins_ × N_cells_, where rows correspond to genomic bins and columns represent individual sampled cells. The copy number state table is utilized for downstream SCNA pattern analysis. To isolate a specific genomic locus, we select the bin segments spanning the cytoband of interest; precisely, the physical start and end positions of the cytoband are mapped to their corresponding boundary bins.

#### Copy number comparison between high- and low-fitness cell groups

The high- and low-fitness cell groups were identified using the base R *kmeans* function (version 4.4.0) applied to the inferred relative cellular fitness values, with the number of cluster centers set differently for different patients to capture the highest fitness group (Figure S3 and Fig-ure S4). We then used the non-parametric Wilcoxon rank-sum test (base R function *wilcox.test*) to compare copy number values at each 500kb bin between the two cell groups. The resulting p-values across all 6087 bins were adjusted for multiple testing using Benjamini-Hochberg pro-cedure (base R function *p.adjust*). The direction of the copy number difference was determined by the sign of the mean difference between groups. We further verified that this direction was consistent with that obtained using the median difference; discrepancies were observed only in bins without significant copy number divergence between groups.

#### Identification of recurrent fitness-associated CN regions

The CN landscape of fitness-associated alterations is characterized by significant inter-patient heterogeneity. Within a single tumor, adjacent genomic bins can exhibit discordant directions of CN change between high- and low-fitness cells. Across patients, a specific region may show significant increased CNs in high-fitness cells in some individuals, but display reduced CNs in others.

To isolate convergent evolutionary signals, we developed a filtering approach to identify recurrent chromosomal regions. The objective of this method is to identify large, contiguous genomic intervals that demonstrate a uniform, statistically significant direction of CN divergence across multiple independent patient tumors. By requiring geonmic contiguity, cross-patient consistency and recurrence, this analysis prioritizes robust signals over patient-specific or localized CN patterns. These regions were identified according to the following criteria:

- **Consistency:** The region shows a uniform direction of statistically significant (p < 0.01) CN divergence between high- versus low-fitness cells across at least 30 consecutive genomic bins (500 kb each). Regions exhibiting heterogeneous directions across patients, demonstrating both increased and decreased CN in high-fitness cells, are filtered out.
- **Recurrence:** The region passes the significance threshold (p < 0.01) in at least three independent patients.

## Data and Code Availability

All patient datasets analyzed in this study were obtained from a previously published study^44^ and accessed through Synapse (access number: syn66366718). The MEDICC2 tree outputs are available on Synapse (access number: syn66481693). The LeafRank package and tutorial are available at GitHub: https://github.com/SunPathLab/LeafRank.

## Acknowledgements

Research in the Sun Lab is supported by NIH grant R01CA276666. This work was supported in part by NSF under Grant No. 2526511. This study uses the computing resources in Minnesota Supercomputing Institute. Zicheng Wang is supported in part by the Key Program of the National Natural Science Foundation of China (NSFC) under Grant No. 72495131 and the Guangdong (China) Provincial Key Laboratory of Mathematical Foundations for Artificial Intelligence [2023B1212010001]. We thank Dr. Anna Selmecki and Xuanming Zhang for helpful discussions.

## Author Contributions

Z.W., C.W., and R.S. designed the study. Z.W., C.W., and R.S. developed the algorithms. Z.W. and C.W. constructed mathematical models. C.W., Z.W., and R.S. performed simulation studies. C.W. and R.S. performed the analysis of single-cell data and visualized the results. C.W., Z.W., K.L., and R.S. interpreted the results and wrote the manuscript. All authors reviewed and provided feedback on the manuscript.

