## Supplemental text and figures for "LeafRank: A phylodynamic framework for inferring relative fitness from single-cell phylogenies in chromosomally unstable tumors"

### Contents

**Notation for the phylogenetic tree and the multitype branching process model** **2**

**Computation of the probability density of a phylogenetic tree** **3**

**Efficient computation of marginal fitness probabilities using a message-passing algorithm** **6**

***In silico* investigation of the effect of birth-rate settings on inference performance** **9**

**Conversion of distance-matrix based phylogenetic trees into ultrametric trees** **11**

**Plotting procedure for the hanging rootogram** **12**

**Supplementary analysis of Truncal WGD patient data** **13**
Cross-patient SCNA analysis reveals fitness-associated chromosomal alterations . . 17

**Supplementary analysis of sub-clonal WGD tumors** **17**
Robustness of OV-075 data analysis under alternative model assumptions . . . . . . 19

| Term | Description |
|---|---|
| Fitness | Cellular net growth rates (difference between birth and death rates). |
| SCNA | Somatic copy number aberration |
| SSNV | Somatic single nucleotide variant |
| abberation | Genomic alterations such as SCNAs and SSNVs in cancer. |
| CN | Copy number |
| CIN | Chromosomal instability |
| $r_S$ | Spearman correlation |
| TT tree | True time ultrametric tree |
| DM tree | Distance-matrix based tree |
| WGD | Whole genome duplication |
| WGD tree | Reconstructed ultrametric tree for a population with subclonal WGD. |
| Truncal WGD | Clone with single WGD event ancestral to all cells, with no residual $0\times$WGD cells. |
| Parallel WGD | Multiple clones with different ancestral WGD events. |
| Subclonal WGD | WGD clone coexisting with $0\times$WGD cells. |
| MRCA | Most recent common ancestor. |

**Table S1.** Terminology and Descriptions.

### Notation for the phylogenetic tree and the multitype branching process model

LeafRank infers relative fitness, defined as the difference between birth and death rates, from single-cell phylogenetic trees using a multitype branching process model. Before describing the implementation of LeafRank, we first introduce the notation used throughout the paper.

Given a single-cell phylogenetic tree, as illustrated in Figure S1A, we distinguish between the set of internal nodes, $\mathcal{Y} = \{y_1, \ldots, y_{N-1}\}$, and the set of leaf nodes, $\mathcal{X} = \{x_1, \ldots, x_N\}$. Recall that, for a binary phylogenetic tree, the number of internal nodes is one less than the number of leaf nodes. We denote the root node by $y_0$. For notational convenience, the left and right immediate descendants of an internal node $y_i$ are denoted by $y_{i,l}$ and $y_{i,r}$, respectively. The ideal input for LeafRank is an ultrametric tree, in which branch lengths represent elapsed time. Accordingly, we denote the bifurcation times along the tree by

$$T = s_0 > s_1 > \cdots > s_{N-1} > 0,$$

as shown in Figure S1A.

In the multitype branching process model, we assume that there are V cell types, where type i is characterized by birth rate $b_i$ and death rate $d_i$. For simplicity, we assume that $b_{i+1} > b_i$, while the death rate is constant across types. Each cell of type i can produce a cell of type $i+1$ at rate $\nu$, which we refer to as the driver aberration rate. We denote by $\mathbf{v}_X = (v_{x_1}, \ldots, v_{x_N})$ and $\mathbf{v}_Y = (v_{y_0}, v_{y_1}, \ldots, v_{y_{N-1}})$ the cell types associated with the leaves and internal nodes, respectively. The root type $v_{y_0}$ is assumed to be the lowest-fitness type; that is,

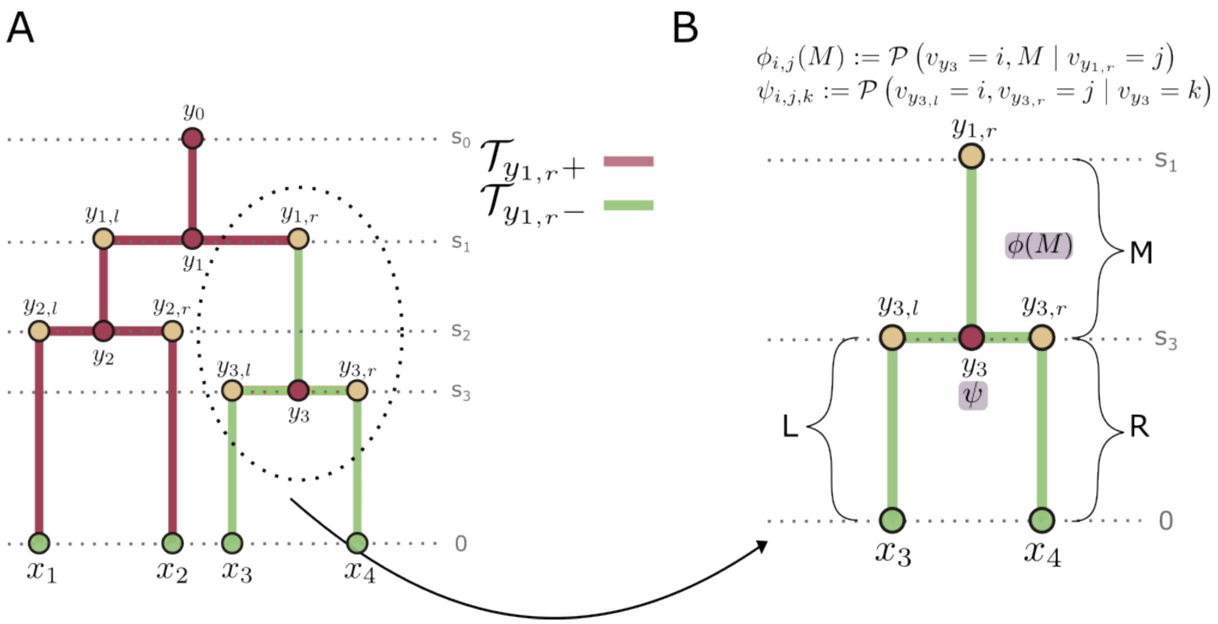


**Figure S1. Message-passing algorithm for efficient computation of marginal probabilities. (A)** Illustrative ultrametric tree. The subtree below ($\mathcal{T}_{y_{1,r}-}$) and the subtree above ($\mathcal{T}_{y_{1,r}+}$) the internal node $y_{1,r}$ are color-coded, respectively. **(B)** Illustrative example of the **Branch Propagator** $\phi$(M) and the **Bifurcation Propagator**.

$$\mathbb{P}(v_{y_0} = 1) = 1.$$

This model can be extended to allow more general type transitions, in which a cell of type i may produce a cell of any type $j \neq i$, for all $j \in \{1, \ldots, V\}$.

Finally, we recall several additional notations. Let $\rho \ll 1$ denote the small probability that a living cell is sampled, and let $\mu$ denote the neutral aberration rate.

### Computation of the probability density of a phylogenetic tree

In this section, we demonstrate the calculation of the density of a phylogenetic tree under the multitype branching process model, following the method proposed in[65]. Recall that $D_{n,v}(t)$ is defined as the density that a lineage starting at time t in state v would evolve into the clade observed to descend from node n. In particular, $D_{y_0,v_{y_0}}(T)$ represents the density that a root node of type $v_{y_0}$ generates the entire observed tree. To evaluate this density at the root, we proceed backward in time, starting from the terminal branches connected to the leaves. The computation relies on two types of propagators: branch propagators and bifurcation propagators. An illustrative computation based on the tree shown in Figure S1B is provided to demonstrate this procedure.

**Calculating** $D_{n,v}(t)$ **along a branch** Consider a leaf $x_3$ in the tree of type $v_{x_3}$. It is evident that $D_{y_{3,l},i}(0) = \rho$ for $i = v_{x_3}$, and $D_{y_{3,l},i}(0) = 0$ for $i \neq v_{x_3}$. Next, we derive a system of differential equations governing $D_{y_{3,l},i}(t)$ for $t > 0$.

Using the first-step method, we express $D_{y_{3,l},i}(t+\Delta t)$ in terms of $D_{y_{3,l},i}(t)$, where $\Delta t$ represents an infinitesimal time step. During the time interval $\Delta t$ along branch L, one of the following events may occur: no event, death, same-type birth, or driver aberration.

$$
\begin{aligned}
D_{y_{3,l},i}(t+\Delta t) = \big(1-(b_i+d_i+\nu)\Delta t\big)\, & D_{y_{3,l},i}(t) \\
& + 2b_i\Delta t E_i(t) D_{y_{3,l},i}(t) \\
& + \nu\Delta t E_{i+1}(t) D_{y_{3,l},i}(t) \\
& + \nu\Delta t E_i(t) D_{y_{3,l},i+1}(t) \\
& + O(\Delta t^2),
\end{aligned}
$$

where $E_i(t)$ is defined as the probability that an individual type i cell has no sampled descendants after time t. Note that when i = V, the driver aberration terms with $\nu$ will disappear. Letting $\Delta t \to 0$, we obtain

$$
\begin{aligned}
\frac{d}{dt}D_{y_{3,l},i}(t) = & -(b_i+\nu+d_i)D_{y_{3,l},i}(t) \\
& + 2b_iE_i(t)D_{y_{3,l},i}(t) \\
& + \nu E_{i+1}(t)D_{y_{3,l},i}(t) \\
& + \nu E_i(t)D_{y_{3,l},i+1}(t).
\end{aligned}
$$

We note that this ODE system is linear as $E_i(t)$ does not contain $D_{y_{3,l},i}(t)$, allowing implementation of principle of superposition. To solve this system of differential equations, we next derive the computation of $E_i(t)$.

**Calculating** $E_i(t)$**.** In this part, we derive a system of differential equations for $E_i(t)$. At time t = 0, the probability that a type i cell is not sampled is given by $1-\rho$, resulting in $E_i(0) = 1-\rho$. The differential equations for $E_i(t)$ are derived using a methodology analogous to the approach outlined earlier for $D_{y_{3,l},i}(t)$.

$$
\begin{aligned}
E_i(t+\Delta t) = & (1-(b_i+\nu+d_i)\Delta t)E_i(t) \\
& + d_i\Delta t \\
& + b_i\Delta t E_i(t)E_i(t) \\
& + \nu\Delta t E_{i+1}(t)E_i(t) \\
& + O(\Delta t^2).
\end{aligned}
$$

Letting $\Delta t \to 0$, we obtain

$$
\begin{aligned}
\frac{d}{dt}E_i(t) = & -(b_i+\nu+d_i)E_i(t) \\
& + d_i \\
& + b_iE_i(t)E_i(t) \\
& + \nu E_{i+1}(t)E_i(t).
\end{aligned}
$$

By solving the system of differential equations for $E_i(t)$ and $D_{y_{3,l},i}(t)$ using the initial values mentioned above, we can calculate $D_{y_{3,l},i}(t)$ along branch L.

**Pruning two subtrees.** We now derive the probability density $D_{y_3,v_{y_3}}(s_3)$ for observing the two sub-trees generated by the bifurcation event at node $y_3$ (of type $v_{y_3}$) occurring at time $s_3$. These sub-trees originate from the left and right immediate children of $y_3$, respectively. While $D_{y_3,v_{y_3}}(s_3)$, $D_{y_{3,l},v_{y_{3,l}}}(s_3)$, and $D_{y_{3,r},v_{y_{3,r}}}(s_3)$ are all computed at time $s_3$, they are distinct quantities at different nodes in the tree connected by the bifurcation event at $y_3$. Let L and R denote the branches descending from $y_3$, belonging to the left and right sub-trees (see Figure S1). Conditional on a bifurcation at node $y_3$, the two immediate descendants generate the left and right observed subtrees. The contribution of this bifurcation event can be written as

$$D_{y_3,k}(s_3) = \sum_{i=1}^{V}\sum_{j=1}^{V} \psi_{i,j,k} D_{y_{3,l},i}(s_3) D_{y_{3,r},j}(s_3),$$

where

$$\psi_{i,j,k} = 2b_k \mathbb{1}_{\{i=j=k\}} + \nu\mathbb{1}_{\{i=k,\ j=k+1\}} + \nu\mathbb{1}_{\{i=k+1,\ j=k\}}.$$

Here, $\psi_{i,j,k}$ is the **Bifurcation Propagator**, giving the rate contribution for a parent of type k to generate left and right descendants of types i and j.

**Decomposition into two propagator functions** Based on the above analysis, the density of a phylogenetic tree can be computed by solving the governing differential equations and recursively pruning subtrees backward in time until $t = s_0$. This procedure decomposes into independent computations over each branch and bifurcation event in the tree. Such a decomposition is particularly advantageous for efficiently computing the marginal distributions of leaf states across the tree.

To illustrate the decomposition method, we consider three branches: M, L and R. Branches L and R are terminal branches, connecting to leaves $x_3$ and $x_4$, respectively, and descend from branch M at the bifurcation event at node $y_3$. Without loss of generality, we assume that branch M originates from the right child of an earlier bifurcation event at node $y_1$, whose state is denoted by $v_{y_{1,r}}$. The probability density $D_{y_{1,r},v_{y_{1,r}}}(s_1)$ is computed in three steps:

I. **Branch L and R:** compute $D_{y_{3,l},v_{y_{3,l}}}(s_3)$ and $D_{y_{3,r},v_{y_{3,r}}}(s_3)$.

II. **Bifurcation:** compute $D_{y_3,v_{y_3}}(s_3)$.

III. **Branch M:** compute $D_{y_{1,r},v_{y_{1,r}}}(s_1)$ using $D_{y_{1,r},i}(s_3) = D_{y_3,i}(s_3)$.

Next, we observe that along each branch, the system of differential equations governing $D_{n,v}(t)$ is linear and time-varying. Consequently, the solution for each branch can be computed independently using basis vectors as initial conditions. Specifically, for branch M, we define a $V \times V$ matrix, called the **Branch Propagator**, whose (i, j)-th entry $\phi_{i,j}(M)$ is zero whenever $j > i$. For $j \leq i$, $\phi_{i,j}(M)$ is obtained by solving for $D_{y_{1,r},v_{y_{1,r}}}(s_1)$ given $v_{y_{1,r}} = j$, $v_{y_3} = i$, and the initial condition: $D_{y_{1,r},i}(s_3) = 1$ and $D_{y_{1,r},j}(s_3) = 0$ for $j \neq i$.

This approach allows us to decompose the computation of $D_{y_{1,r},v_{y_{1,r}}}(s_1)$ into three independent parallel components: calculating the density vectors $\{D_{y_{3,l},i}(s_3)\}_{i=1}^{V}$ and $\{D_{y_{3,r},j}(s_3)\}_{j=1}^{V}$, and calculating the branch propagator $\phi_{i,j}(M)$ for $i, j \in \{1, \ldots, V\}$. These components are first combined through the bifurcation propagator $\psi_{i,j,k}$ at node $y_3$:

$$D_{y_3,k}(s_3) = \sum_{i=1}^{V}\sum_{j=1}^{V} \psi_{i,j,k} D_{y_{3,l},i}(s_3) D_{y_{3,r},j}(s_3), \qquad k = 1, \ldots, V.$$

The resulting density vector at node $y_3$ is then propagated along branch M from time $s_3$ to time $s_1$. Under the convention that $\phi_{k,j}(M)$ maps descendant type k at node $y_3$ to ancestral type j at node $y_{1,r}$, we obtain (by the Principle of Superposition)

$$D_{y_{1,r},j}(s_1) = \sum_{k=1}^{V} \phi_{k,j}(M) D_{y_3,k}(s_3).$$

Therefore, for the specific state $v_{y_{1,r}}$ at node $y_{1,r}$,

$$D_{y_{1,r},v_{y_{1,r}}}(s_1) = \sum_{k=1}^{V} \phi_{k,v_{y_{1,r}}}(M) D_{y_3,k}(s_3).$$

Substituting the bifurcation-propagator expression for $D_{y_3,k}(s_3)$ gives

$$D_{y_{1,r},v_{y_{1,r}}}(s_1) = \sum_{k=1}^{V}\sum_{i=1}^{V}\sum_{j=1}^{V} \phi_{k,v_{y_{1,r}}}(M)\psi_{i,j,k} D_{y_{3,l},i}(s_3) D_{y_{3,r},j}(s_3).$$

Thus, the density at node $y_{1,r}$ can be computed by independently evaluating the two descendant subtree density vectors, combining them with the bifurcation propagator at $y_3$, and then applying the branch propagator along branch M. The same argument applies when branches L and R subtend internal subtrees rather than leaves, because each subtree can be summarized by a density vector at its root. Recursively applying the branch and bifurcation propagators from the leaves to the root yields $D_{y_0,v_{y_0}}(T)$, the density of the full observed phylogenetic tree.

### Efficient computation of marginal fitness probabilities using a message-passing algorithm

For a given tree $\mathcal{T}$, the goal of LeafRank is to compute the marginal probability of the fitness type at each leaf, conditioned on the tree topology, i.e.

$$\mathcal{P}\left(v_i \mid \mathcal{T}\right), \quad i \in \mathcal{X}.$$

As mentioned in Method, directly summing over all possible joint assignments of node types is computationally infeasible as the number of fitness type and the tree size increase. In this section, we present an efficient strategy for computing these marginal probabilities at each leaf using a message passing algorithm[64].

We first denote by $\mathcal{T}_{y_i-}$ and $\mathcal{T}_{y_i+}$ the subtrees below and above the node $y_i$, respectively. Notably, $\mathcal{T}_{y_i+}$ contains all the information upstream of $y_i$, including contributions from its sibling subtree. Using this notation, we define two key conditional probabilities/densities, which serve as messages in our message passing algorithm:

- $\mathcal{P}\left(\mathcal{T}_{y_i-} \mid v_{y_i}\right)$: The density of generating the tree below node $y_i$ given node $y_i$'s type.
- $\mathcal{P}\left(v_{y_i} \mid \mathcal{T}_{y_i+}\right)$: The probability mass of node $y_i$'s type given the ancestry and sibling evolutionary information $\mathcal{T}_{y_i+}$.

The marginal probability of node $y_i$'s type then follows from these two quantities:

$$\mathcal{P}\left(v_{y_i} \mid \mathcal{T}\right) \propto \mathcal{P}\left(\mathcal{T}_{y_i-} \mid v_{y_i}\right) \mathcal{P}\left(v_{y_i} \mid \mathcal{T}_{y_i+}\right).$$

To show this relationship, we introduce an important observation from the classic Felsenstein likelihood work[70]:

**Observation 1.** $\mathcal{P}\left(\mathcal{T}_{y_i-} \mid v_{y_i}\right) = \mathcal{P}\left(\mathcal{T}_{y_i-} \mid v_{y_i}, \mathcal{T}_{y_i+}\right)$ *due to conditional independence between* $\mathcal{T}_{y_i-}$ *and* $\mathcal{T}_{y_i+}$ *given* $v_{y_i}$*.*

Next, we recall that the branch propagator $\phi_{ij}(L)$ is proportional to $\mathcal{P}\left(v_{x_3} = i, L \mid v_{y_{3,l}} = j\right)$. With this and bifurcation propagator, we have:

- **Branch Propagator:**

$$\begin{aligned}
&\mathcal{P}\left(\mathcal{T}_{y_{3,l}-} \mid v_{y_{3,l}} = j\right) = \mathcal{P}\left(\mathcal{T}_{x_3-}, L \mid v_{y_{3,l}} = j\right) \\
&= \sum_{i=1}^{V} \mathcal{P}\left(\mathcal{T}_{x_3-}, v_{x_3} = i, L \mid v_{y_{3,l}} = j\right) \\
&= \sum_{i=1}^{V} \mathcal{P}\left(\mathcal{T}_{x_3-} \mid v_{x_3} = i, L, v_{y_{3,l}} = j\right) \mathcal{P}\left(v_{x_3} = i, L \mid v_{y_{3,l}} = j\right) \\
&\propto \sum_{i=1}^{V} \mathcal{P}\left(\mathcal{T}_{x_3-} \mid v_{x_3} = i\right) \phi_{ij}(L).
\end{aligned}$$

- **Bifurcation Propagator:**

$$\begin{aligned}
&\mathcal{P}\left(\mathcal{T}_{y_3-} \mid v_{y_3} = i\right) \\
&= \sum_{j,k} \mathcal{P}\left(\mathcal{T}_{y_{3,l}-}, \mathcal{T}_{y_{3,r}-}, v_{y_{3,l}} = j, v_{y_{3,r}} = k \mid v_{y_3} = i\right) \\
&= \sum_{j,k} \mathcal{P}\left(\mathcal{T}_{y_{3,l}-} \mid v_{y_{3,l}} = j\right) \mathcal{P}\left(\mathcal{T}_{y_{3,r}-} \mid v_{y_{3,r}} = k\right) \mathcal{P}\left(v_{y_{3,l}} = j, v_{y_{3,r}} = k \mid v_{y_3} = i\right). \\
&= \sum_{j,k} \underbrace{\mathcal{P}\left(\mathcal{T}_{y_{3,l}-} \mid v_{y_{3,l}} = j\right)}_{\text{up message from left child}} \underbrace{\mathcal{P}\left(\mathcal{T}_{y_{3,r}-} \mid v_{y_{3,r}} = k\right)}_{\text{up message from right child}} \psi_{j,k,i}
\end{aligned}$$

**Remark 1.** $\mathcal{T}_{y_{3,l}-}$ *is independent of* $\mathcal{T}_{y_{3,r}-}$ *given the types* $v_{y_{3,l}}$ *and* $v_{y_{3,r}}$*.*

We then denote $\mathcal{P}\left(\mathcal{T}_{n-} \mid v_n\right)$ as the "**up message**", which can be compute recursively from leaf nodes to root nodes.

For another quantity, $\mathcal{P}\left(v_{y_i} \mid \mathcal{T}_{y_i+}\right)$ denoted as "**down message**", we start from the root of tree. In particular, we assume that the root fitness type is the lowest type, $\mathcal{P}\left(v_{root} = 1\right) = 1$, which is the only assumption we made about the node fitness type. In other words, all the relative fitness obtained from LeafRank based on this assumption and we basically derived the fitness at leaves relative to the root fitness type. The probability mass of root's type is then propagate along the tree to all the leaves based on

- **Branch Propagator:**

$$\begin{aligned}
\mathcal{P}\left(v_{y_3} = j \mid \mathcal{T}_{y_3+}\right) &= \sum_i \mathcal{P}\left(v_{y_3} = j, v_{y_{1,r}} = i \mid M, \mathcal{T}_{y_{1,r}+}\right) \\
&\propto \sum_i \mathcal{P}\left(v_{y_3} = j, v_{y_{1,r}} = i, M \mid \mathcal{T}_{y_{1,r}+}\right) \\
&= \sum_i \mathcal{P}\left(v_{y_3} = j, M \mid v_{y_{1,r}} = i\right) \mathcal{P}\left(v_{y_{1,r}} = i \mid \mathcal{T}_{y_{1,r}+}\right) \\
&\propto \sum_i \phi_{ji}(M) \mathcal{P}\left(v_{y_{1,r}} = i \mid \mathcal{T}_{y_{1,r}+}\right).
\end{aligned}$$

- **Bifurcation Propagator:**

$$\begin{aligned}
&\mathcal{P}\left(v_{y_{3,l}} = i \mid \mathcal{T}_{y_{3,l}+}\right) \\
&= \sum_{j,k} \mathcal{P}\left(v_{y_{3,l}} = i, v_{y_{3,r}} = j, v_{y_3} = k \mid \mathcal{T}_{y_3+}, \mathcal{T}_{y_{3,r}-}\right) \\
&\propto \sum_{j,k} \mathcal{P}\left(\mathcal{T}_{y_{3,r}-}, v_{y_{3,l}} = i, v_{y_{3,r}} = j, v_{y_3} = k \mid \mathcal{T}_{y_3+}\right) \\
&= \sum_{j,k} \mathcal{P}\left(\mathcal{T}_{y_{3,r}-} \mid v_{y_{3,r}} = j\right) \mathcal{P}\left(v_{y_{3,l}} = i, v_{y_{3,r}} = j, v_{y_3} = k \mid \mathcal{T}_{y_3+}\right) \\
&= \sum_{j,k} \mathcal{P}\left(\mathcal{T}_{y_{3,r}-} \mid v_{y_{3,r}} = j\right) \mathcal{P}\left(v_{y_{3,l}} = i, v_{y_{3,r}} = j \mid v_{y_3} = k\right) \mathcal{P}\left(v_{y_3} = k \mid \mathcal{T}_{y_3+}\right) \\
&= \sum_{j,k} \underbrace{\mathcal{P}\left(\mathcal{T}_{y_{3,r}-} \mid v_{y_{3,r}} = j\right)}_{\text{up message from sibling}} \psi_{i,j,k} \underbrace{\mathcal{P}\left(v_{y_3} = k \mid \mathcal{T}_{y_3+}\right)}_{\text{down message from ancestor}}
\end{aligned}$$

**Remark 2.** *This bifurcation update combines the down message from the ancestor with the up message from the sibling subtree.*

These computations efficiently propagate information from the root to each node under the root-type assumption. Once the down message has been propagated to the leaves, we can see $\mathcal{P}\left(v_{x_3} \mid \mathcal{T}_{x_3+}\right)$ is exactly the marginal probability for leaf node $x_3$ conditioned on the whole tree.

### *In silico* investigation of the effect of birth-rate settings on inference performance

In the main Results, we investigated the effects of parameter perturbations on fitness ranking and identified two structural patterns in the inferred fitness densities that serve as explicit indicators of a noninformative fitness prediction. Across all experiments, only the b1 and b2 settings exhibited both uninformative patterns and a noticeable decline in accuracy across 50 virtual tumor experiments. In these settings, the number of fitness states was reduced by modifying the birth-rate settings while keeping all other input parameters fixed. Motivated by this observation, we provide a detailed analysis of how changes in the birth-rate settings, which parameterize the underlying fitness states, influence the inference results.

In the b1 and b2 settings, we hypothesized that using fewer assumed fitness states limits the model's capacity to accurately infer rankings among sampled leaves. In the extreme case where only a single fitness state is assumed, all leaves are assigned to the same state, and no meaningful ranking can be inferred. Although both configurations exhibited degraded performance relative to the original parameter configurations, b1, which uses the same logarithmic spacing scheme as the underlying simulation, achieves more accurate inference than b2, which uses a linear spacing scheme. We therefore hypothesized that matching the assumed and underlying parameter spacing schemes improves ranking inference accuracy. These two spacing schemes correspond to different assumptions about driver aberration effects: a logarithmic configuration implies multiplicative fitness gains, whereas a linear configuration assumes additive increases. To test this hypothesis, we conducted two-parameter sensitivity analysis experiments using virtual tumors simulated under both linear and logarithmic spacing schemes.

In particular, we simulated a new set of *in silico* tumors (*T-lin*, with 50 counts) under the non-spatial simulation framework, in which driver aberrations increased birth rate linearly from 0.2 to 0.8. Using these newly generated *T-lin* tumors, together with the original *in silico* tumor (*T-log*) characterized by logarithmic spacing, we evaluated inference performance under the parameter configurations listed in Table S2. The resulting fitness ranking accuracies for *T-lin* and *T-log* are presented in Figure S2A and Figure S2B, respectively.

| | Time Scale $\tau$ = 1 | Time Scale $\tau$ = 2 | Time Scale $\tau$ = 4 |
|---|---|---|---|
| $\mathcal{C}1 : b_i \in \{0.2 \cdot 1.2^i\}_{i=0,\cdots,7}$ | $\tau 1\mathcal{C}1$ | $\tau 2\mathcal{C}1$ | $\tau 4\mathcal{C}1$ |
| $\mathcal{C}2 : b_i \in \{0.2 + 1.2 \cdot i\}_{i=0,\cdots,7}$ | $\tau 1\mathcal{C}2$ | $\tau 2\mathcal{C}2$ | $\tau 4\mathcal{C}2$ |
| $b1 : b_i \in \{0.2, 0.31, 0.47, 0.72\}$ | $\tau$1b1 | $\tau$2b1 | $\tau$4b1 |
| $b2 : b_i \in \{0.2, 0.37, 0.54, 0.72\}$ | $\tau$1b2 | $\tau$2b2 | $\tau$4b2 |

**Table S2.** Two-parameters sensitivity analysis configurations. The configuration $\mathcal{C}$ use the ground-truth fitness types during the simulation.

We found that correctly specifying the fitness phenotype spacing scheme did not necessarily yield more accurate ranking inference. For example, in analyses based on *T-log* tumors, the $\tau$4b2 configuration, which assumes a linear spacing scheme, achieved slightly better performance than the configuration $\tau 4\mathcal{C}1$. Similarly, in analyses bsed on *T-lin* tumors, the $\tau$2b1 configuration achieved inference accuracy comparable to that of $\tau 2\mathcal{C}2$. These results demonstrate that ranking inference accuracy is decoupled from the precise configuration of the underlying fitness phenotype spacing scheme.

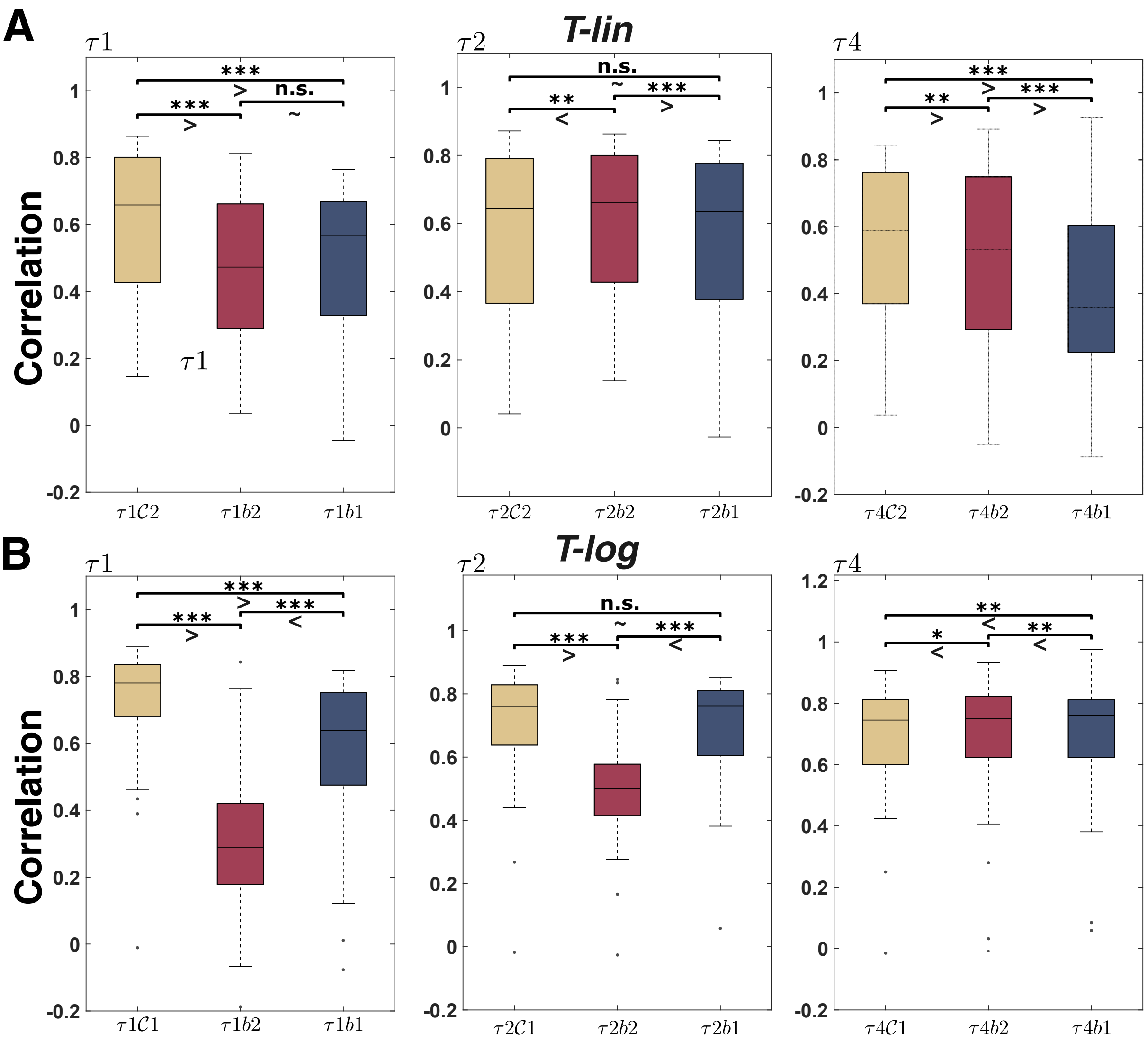


**Figure S2.** Two-parameter sensitivity analysis on *in silico* tumor *T-lin* and *T-log*. **(A)** Inferred ranking accuracy across 50 *T-lin in silico* tumors under the corresponding configurations listed in Table S2. Statistical significance was assessed using the Wilcoxon signed-rank test (n.s.: $p \geq 0.05$, *: $p < 0.05$, **: $p < 0.01$, ***: $p < 0.001$). The dominant ordinal direction between each pair of groups is shown below the corresponding significance annotation. **(B)** Inferred ranking accuracy across 50 *T-log in silico* tumors under the corresponding configurations in listed in Table S2. Arrangement and notation are the same as in (A).

To further investigate the impact of fitness spacing schemes, we compared the inferred fitness distributions in cases where b1 outperforms b2 and vice versa. Increasing the time scale factor enlarges the inferred leaf fitness values by shortening the elapsed evolutionary time between the MRCA and the sampled leaves. When b1 performed better, as observed for T-log under $\tau$1 and $\tau$2, the majority of inferred leaf fitness values were concentrated in the lower fitness range. Conversely, when b2 performs better as observed for T-lin under $\tau$2 and $\tau$4, the inferred fitness values were more concentrated in the higher fitness range. These performance differences can be interpreted through the resolution induced by each fitness spacing scheme. Logarithmic spacing places fitness states more densely in the lower fitness range and more sparsely in the higher fitness range, allowing the $\tau$1b1 and $\tau$2b1 configurations to better resolve variation when inferred leaf fitness values are concentrated at lower values. In contrast, linear spacing distributes fitness states uniformly across the full range, providing relatively grater resolution in the higher fitness regime and thereby improving sensitivity when inferred leaf fitness values are concentrated at higher values.

Together, these results suggest that selecting informative parameter configurations should provide sufficient fitness-state resolution in the regions where the inferred fitness density is concentrated. In the Methods section, we provide a systematic approach for identifying empirically feasible configurations that satisfy this requirement.

### Conversion of distance-matrix based phylogenetic trees into ultrametric trees

To construct an ultrametric tree from a distance-matrix based phylogenetic tree, we adapted the molecular dating algorithm implemented in *chronos*[46]. The *chronos* function estimates a chronogram in which branch lengths represent the temporal distance between ancestor and descendant nodes. The method supports several evolutionary models, including *correlated*, *discrete*, and *strict clock* models, all of which assume that aberrations accumulate according to a Poisson process. Under this assumption, the number of aberrations along branch i, denoted by $X_i$, follows a Poisson distribution with rate parameter $\xi_i$:

$$\mathbb{P}(X_i = x_i | \xi_i) = \xi_i^{x_i} \frac{e^{-\xi_i}}{x_i!}.$$

The expected number of aberrations on branch i, denoted by $\xi_i := r_i t_i$, is typically modeled as the product of aberration rate $r_i$ and temporal duration $t_i$ of the branch. In standard molecular dating formulations, the aberration rate is often assumed to be constant across branches, such that $r_i = r$. Under this assumption, a maximum likelihood estimation (MLE) framework can be used to estimate the parameter set $(r, \{t_i\}_{i \in \mathbf{B}})$, where $\mathbf{B}$ denotes the set of all branches. Specifically, the parameters are estimated by minimizing the negative log-likelihood:

$$\min_{r, \{t_i\}_{i \in \mathbf{B}}} -\left( \sum_{i \in \mathbf{B}} x_i \log(r t_i) - r t_i - \log(x_i!) \right),$$

where terms independent of the parameters have been omitted. For phylogenetic trees reconstructed from samples collected at a single time point, an ultrametric constraint is imposed such that $\sum_{i \in \mathbf{B}_\ell} t_i = 1$, for every leaf-to-root path $\mathbf{B}_\ell$, ensuring that all leaves are equidistant from the root in inferred temporal distance.

However, the current implementation of *chronos* does not account for phylogenetic trees containing both genome doubled ($WGD^+$) and diploid ($WGD^-$) lineages. In this setting, the elevated chromosomal instability associated with $WGD^+$ is expected to induce substantially higher aberration rates[39–41]. To accommodate this biological heterogeneity, we extended the original *chronos* framework to allow aberration-rate shifts associated with WGD events. This extension permits state-dependent aberration rates and provides a more biologically realistic temporal calibration for phylogenetic trees reconstructed from copy number profiles.

Specifically, we first classify each branch according to the WGD status of its descendant node and make the following assumption.

**Assumption** (*Anc.*)**.** *For each $WGD^+$ clade, the WGD event is assigned to the ancestral end of the branch leading to the earliest node inferred to be $WGD^+$. Consequently, the branch leading to this earliest $WGD^+$ node is labeled as $WGD^+$, together with all branches descending from this node.*

We denote the sets of $WGD^-$ and $WGD^+$ branches by $\mathbf{B}_{WGD-}$ and $\mathbf{B}_{WGD+}$, respectively, and assign distinct aberration rates $r_-$ and $r_+$ to the two branch classes. We note that this assumption produces a branch classification that is largely concordant with the WGD classification inferred in the original study[44]. To accommodate WGD-associated rate shifts, we further introduce the following assumption.

**Assumption** (*Poi.*)**.** *The $WGD^+$ population accumulates SCNAs according to the same Poisson process as the $WGD^-$ population, but with an approximately twofold higher aberration rate.*

To capture the elevated aberration rate associated with WGD, we formulated a constrained maximum likelihood estimation problem by imposing $r_+ \in (1.03r_-, 2.5r_-)$, based on recent empirical studies[40,44]. This constraint reflects the biological expectation that WGD increases the aberration rate while still allowing flexibility in parameter estimation.

Under these assumptions, the optimization problem becomes:

$$\min_{r_-, r_+, \{t_i\}_{i\in\mathbf{B}}} \left( \sum_{i\in\mathbf{B}_{WGD-}} x_i \log(r_- t_i) - r_- t_i \right) + \left( \sum_{i\in\mathbf{B}_{WGD+}} x_i \log(r_+ t_i) - r_+ t_i \right),$$

where terms independent of the parameters have been omitted. This constrained optimization problem was solved in MATLAB using the interior-point algorithm implemented in *fmincon* function[68]. Due to the non-convexity of the optimization problem, we applied a multi-start strategy to identify the optimal solution for ultrametric tree reconstruction.

### Plotting procedure for the hanging rootogram

In the Results, we used hanging rootograms to assess the fit of the reconstructed ultrametric trees to the original phylogenies. Here, we describe the construction of these plots and briefly summarize the underlying rationale. For a more comprehensive rationale, we refer readers to the original study[69].

To assess how well the reconstructed ultrametric tree captures the distribution of copy number aberrations in the original phylogenetic tree under the probabilistic model, we compared the expected (exp) and observed (obs) event frequencies. This comparison is straightforward when the probabilistic model is parameterized by a single shared parameter across all observations. However, in the context of ultrametric tree reconstruction, the inferred temporal durations vary across branches, resulting in branch-specific Poisson parameters. Under the Poisson model, the number of CNAs on branch i is assumed to follow a Poisson distribution with rate $\xi_i = rt_i$. Consequently, rather than treating branches as independent and identical distributed random variable, we considered the aggregate distribution across all branches. The expected frequency of observing j events was computed as

$$\mathrm{exp}_j = \sum_{i \in \mathbf{B}} \mathbb{P}(j; \xi_i),$$

where **B** denotes the set of all branches and $\mathbb{P}(\cdot; \xi_i)$ denotes the probability mass function of a Poisson distribution with parameter $\xi_i$. The hanging rootogram visualizes the expected frequencies as a smooth curve and the observed frequencies as downward-extending bars. Following the original study[69], a square-root transformation was applied to reduce scale differences between frequencies.

### Supplementary analysis of Truncal WGD patient data

In this section, we provide a detailed analysis of the inference results for patients with truncal WGD. Specifically, we examine relative fitness estimates across ten patients, evaluate the validity of the key assumptions, and conclude with a cross-patient analysis of aneuploidy profiles.

#### Relative fitness of patients with truncal WGD suggest two evolutionary patterns with ongoing selection

We begin by presenting detailed inference results for all patients with truncal WGD. Figure S3 and Figure S4 summarize fitness estimates derived from both the original MEDICC2 trees and our converted ultrametric trees, together with the corresponding fitness density plots, the Spearman correlation coefficient ($r_s$) between the two sets of inferences, and hanging rootograms evaluating the ultrametric tree reconstruction. The parameter configurations are reported in Table S3, following the rational described in the Methods.

| Sample | $\rho$ | $b_i$ | $d_i$ | $\nu$ | $\tau$ | $\Delta$ |
|---|---|---|---|---|---|---|
| OV-003 | 0.0005 | $\{1.1 \cdot 1.1^i\}_{i=0,\cdots,15}$ | 1 | 0.0001 | 0.047 | 0.01 |
| OV-008 | 0.0005 | $\{1.1 \cdot 1.1^i\}_{i=0,\cdots,15}$ | 1 | 0.0001 | 0.048 | 0.01 |
| OV-014 | 0.0005 | $\{1.1 \cdot 1.1^i\}_{i=0,\cdots,15}$ | 1 | 0.0001 | 0.045 | 0.01 |
| OV-049 | 0.0005 | $\{1.1 \cdot 1.1^i\}_{i=0,\cdots,15}$ | 1 | 0.0001 | 0.047 | 0.01 |
| OV-068 | 0.0005 | $\{1.1 \cdot 1.1^i\}_{i=0,\cdots,15}$ | 1 | 0.0001 | 0.045 | 0.01 |
| OV-082 | 0.0005 | $\{1.1 \cdot 1.1^i\}_{i=0,\cdots,15}$ | 1 | 0.0001 | 0.044 | 0.01 |
| OV-087 | 0.001 | $\{1.1 \cdot 1.1^i\}_{i=0,\cdots,15}$ | 1 | 0.0001 | 0.063 | 0.01 |
| OV-105 | 0.0005 | $\{1.1 \cdot 1.1^i\}_{i=0,\cdots,15}$ | 1 | 0.0001 | 0.045 | 0.01 |
| OV-118 | 0.0005 | $\{1.1 \cdot 1.1^i\}_{i=0,\cdots,15}$ | 1 | 0.0001 | 0.045 | 0.01 |
| OV-129 | 0.0005 | $\{1.1 \cdot 1.1^i\}_{i=0,\cdots,15}$ | 1 | 0.0001 | 0.043 | 0.01 |

**Table S3.** Summary of inference parameter configurations applied to patients exhibiting Truncal WGD

Fitness estimates obtained from the original MEDICC2 trees and the converted ultrametric trees were highly consistent, with an average Spearman correlation coefficient of $r_s = 0.88$. Two probabilistic models were considered in the ultrametric tree conversion: a Poisson (Poi.) model and a negative binomial (N.b.) model. Owing to the superior performance of Poisson model (in 6 out of 10 patients) and its relative simplicity, the final ultrametric trees were reconstructed using the Poisson model.

Based on the inferred fitness landscapes, we classified patients into two distinct evolutionary patterns: **directional selection** and **parallel selection**. As illustrated in Figure S3, four patients were classified into the **directional selection** category. Consistent with the illustrative example shown in Figure 5A, these tumors exhibited a predominantly unidirectional evolutionary trajectory, in which the highest-fitness sublineage emerged from an already advantaged lineage. At the same time, we consistently observed small low-fitness sublineages that persisted throughout tumor evolution.

Together, these features resemble the classical linear evolution model[5,71], in which tumors acquire driver aberrations in a stepwise manner. Under this model, cells harboring advantageous genomic profiles undergo selective sweeps and progressively dominate the population. Indeed, in patients OV-008 and OV-087, the highest-fitness populations comprises the majority of sampled cells, whereas low-fitness populations were rarely observed.

Nevertheless, LeafRank also identifies intermediate-fitness subpopulations that remained relatively abundant and have not yet been outcompeted, as observed in patient OV-105. These observations suggest that fitness acquisition was an ongoing evolutionary process rather than a completed selective event in these tumors. From a topological perspective, these phylogenies were characterized by an expanded and relatively balanced high-fitness clade near the upper portion of the tree, consistent with the progressive expansion of advantaged lineages.

In contrast, the remaining patients, classified as exhibiting **parallel selection** in Figure S4, showed fitness increases not only within the initially advantaged lineage but also across less advantaged lineages. For example, in patient OV-118, both major clades descending from the root independently gave rise to high-fitness sublineages, resulting in comparable population sizes between the two clades. Within each clade, we additionally observed sublineages with intermediate or low-fitness, further supporting a parallel pattern of evolutionary adaptation.

This pattern is consistent with the branching evolution model proposed in[71], in which multiple lineages independently acquire advantageous genomic alterations. Unlike the **directional selection** pattern, **parallel selection** pattern may involve not only sweeps but also potential cooperation or competition among simultaneously expanding sublineages, raising several intriguing evolutionary questions[72,73]. LeafRank enables the identification and characterization of these sublineages, thereby facilitating further investigation into their interaction, evolutionary dynamics, and underlying biological mechanisms.

Across both evolutionary patterns, we observed not only high-fitness cell groups that comprised the majority of the sampled population, but also low-fitness cell groups that were rarely sampled. These observations suggest that post-WGD evolution was shaped by ongoing selection rather than neutral expansion. To further investigate how this selective process relates to genomic subclonal architecture, we next examine cross-patient patterns of SCNAs.

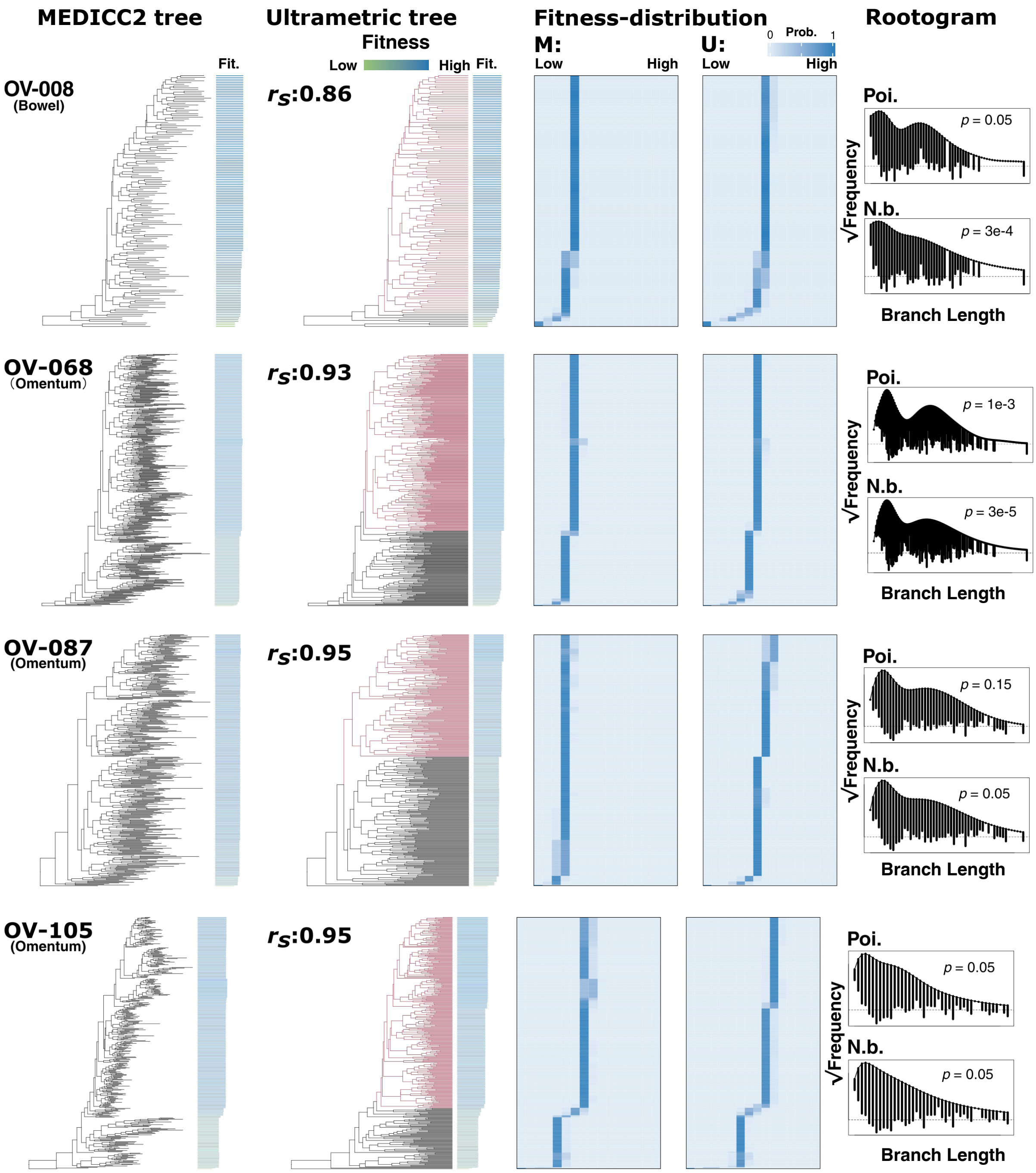


**Figure S3. Fitness rankings in truncal WGD patients reveal directional selection** Each row corresponds to a patient sample with sampling site and contains four subpanels (left to right): **MEDICC2 tree:** the original MEDICC2 tree with inferred fitness estimates, displayed similarly to Figure 5A. **Ultrametric tree:** the reconstructed ultrametric tree with inferred fitness estimates displayed as in MEDICC2 tree. The Spearman rank correlation between fitness rankings inferred from the ultrametric and MEDICC2 trees is reported as $r_S$. Lineages classified as high-fitness are highlighted in red for each patient. **Fitness-distribution:** inferred marginal fitness distributions from the MEDICC2 (M) and ultrametric (U) trees. **Rootogram:** hanging rootograms for ultrametric tree reconstruction under the Poisson (Poi.) and negative binomial (N.b.) models; goodness of fit is evaluated using the Wald-Wolfowitz runs test for randomness of residual signs across branch lengths, with p-values reported in each panel.

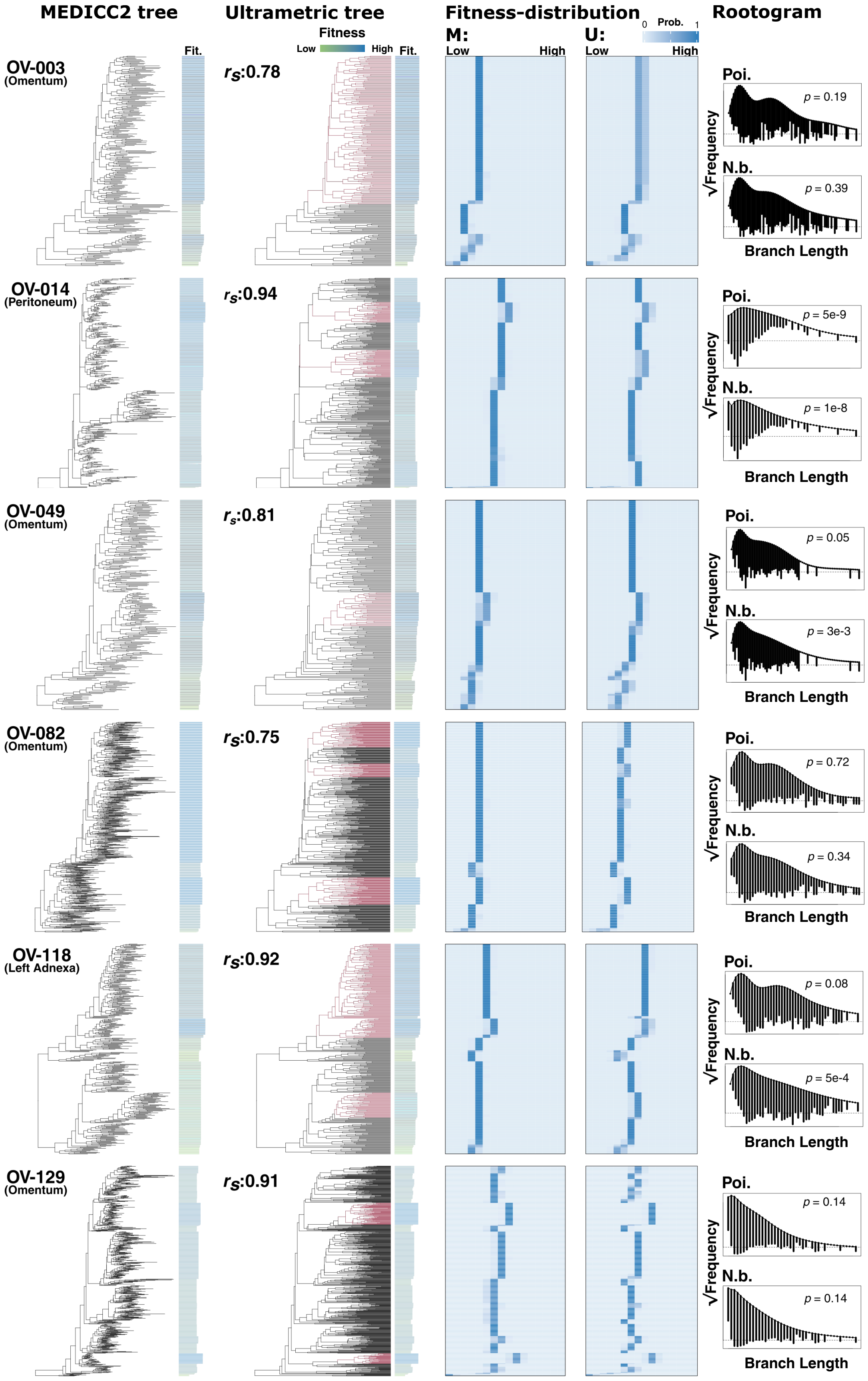


**Figure S4. Fitness Rankings in Truncal WGD patients reveal parallel selection.** Each row represents a patient sample and includes four subpanels, arranged as in Figure S3.

#### Cross-patient SCNA analysis reveals fitness-associated chromosomal alterations

We begin with an overview of SCNA profiles across 10 patients with truncal WGD. Specifically, we examined the average CN relative to the baseline tetraploid state (4 copies) within WGD populations (Figure S5A), a metric analogous to CN measurements derived from bulk sequencing data. Consistent with prior observations[44,56], numerous chromosomal regions exhibited copy numbers below 4. This pattern suggests frequent CN losses occurring either pre-WGD, which can lead to copy-neutral loss of heterozygosity (LOH), or post-WGD, resulting in a localized reversion to triploid states.

We additionally observed recurrent CN imbalances between the p and q arms in chromosome 3 and 8, where the q arms have a CN greater than 4. This is in agreement with prior reports[74,75]. However, the substantial chromosomal instability characteristic of WGD tumors generates highly heterogeneous CN landscapes, limiting the ability of average profiles to resolve fitness-associated SCNA profiles. To address this limitation, we leveraged the fitness estimates inferred by LeafRank and performed a cross-patient analysis of fitness-associated SCNAs by comparing high- and low-fitness cell groups.

Specifically, we applied a K-means clustering approach to stratify single cells into high- and low-fitness groups based on their inferred relative fitness (high-fitness lineages highlighted in Figure S3 and Figure S4). We then compared segment-level CNs between the two groups using Wilcoxon rank-sum tests, with multiple-testing correction performed using the Benjamini-Hochberg procedure. The results are summarized in Figure S5B.

We observed substantial inter-patient heterogeneity, with many genomic segments exhibiting divergent CN trajectories across the high-fitness groups of different patient tumors. Nevertheless, several chromosomal regions displayed recurrent, fitness-associated SCNAs across the cohort. These regions are highlighted in Figure S5B and were selected according to the consistency and recurrence criteria described in the Methods. We hypothesize that these recurrently altered regions harbor key fitness-modulating target genes. For example, the recurrent CN reduction observed on Chr 10 (p11.1 - q24.2) encompasses the well-known tumor suppressor gene *PTEN*. Complementing this loss, we identified a recurrent CN gain on Chr 12 (p13.31 - p11.23) in the high-fitness populations of six patients; this locus harbors the frequently reported oncogene *KRAS*[52,53] (Figure S5C). Furthermore, we detected a recurrent CN reduction on Chr 6 (q15 - q16.3) in the high-fitness groups of six patients (Figure S5C). This region contains *MAP3K7*, a candidate mediator of adaptive cellular signaling; CN depletion of this region has previously been associate with enhanced cellular adaptability under chemotherapy[54]. Beyond these prominent loci, recurrent SCNAs identified on Chr 3, Chr 5, Chr 7, Chr 11, Chr 17, and Chr X represent additional candidate regions potentially driving enhanced cellular fitness in truncal WGD HGSOC (Figure S5B).

Collectively, these findings highlight candidate fitness-associated SCNAs and provide a foundation for further investigation into tumor evolution and selective trajectories inferred from genomic copy-number profiles.

### Supplementary analysis of sub-clonal WGD tumors

Sub-clonal WGD patient datasets (3 patients) represent a uniquely valuable yet complex dataset for evolutionary analysis, providing an ideal benchmark for testing the boundaries of LeafRank.

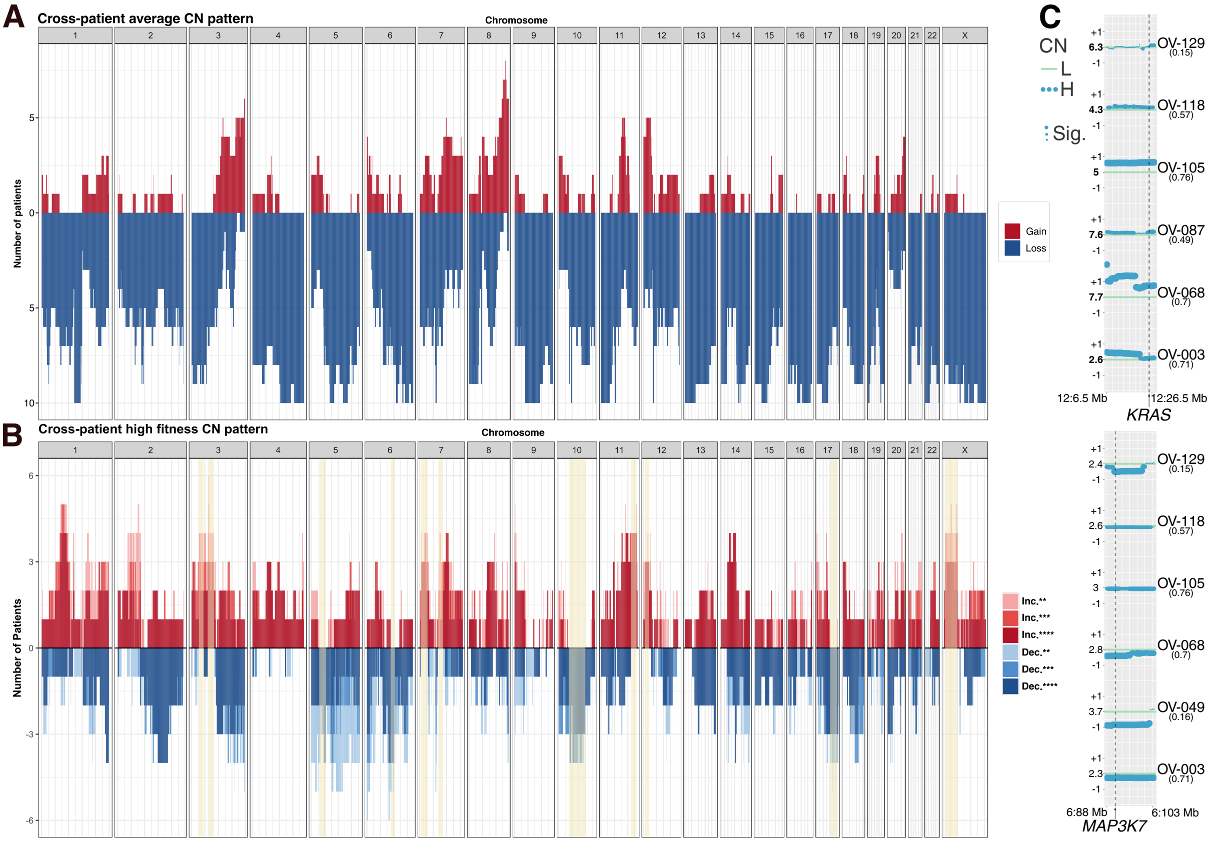


**Figure S5. Cross-patient CNA analysis reveals consistent aneuploidy profiles (A)** Cross-patient average copy number profiles for 10 patients with truncal WGD. The gain and loss are determined by the difference between average copy number and WGD cell baseline ($4 \pm 0.5$). **(B)** Cross-patient high-fitness copy number profiles. The increase (Inc.) and decrease (Dec.) are defined by comparing the high-fitness cells and low-fitness counterparts through Wilcoxon tests. Significant levels above $**$, ($p \leq 0.01$), are included and color-coded differently. The shaded yellow regions highlights the region where increase or decrease signals are consistent and recurrent across patients. **(C)** Highlighted regions on Chr 12 (p13.31 - p11.23) and Chr 6 (q15 - q16.3) identify recurrent, fitness-associated SCNAs across the cohort, formatted as in Figure 6B. The Chr 6 highlighted interval corresponds directly to the region shown in panel (B). In contrast, the Chr 12 region was extended to encompass a locally contiguous signal that was interrupted by one or two isolated genomic bins failing the strict consistency and recurrence criteria detailed in the Methods.

While reconstructing the precise temporal architecture of ultrametric phylogenetic trees, including the exact placement of WGD events, presents an intricate challenge, LeafRank is specifically designed to navigate these data-sparse regimes. To ensure analytical rigor, our framework utilizes principled modeling assumptions to resolve these complex topologies. In this section, we extend our evaluation to additional patient cohorts and demonstrate the robustness of our core conclusions across a wide spectrum of alternative modeling configurations.

### Fitness ranking of WGD$^+$ in other subclonal patient

In Results, we analyzed fitness inference using patient OV-075 data, while two other patient data (OV-006 and OV-139) were also available for analysis. Here, we provide a more detailed analysis of these additional subclonal WGD patient datasets. The corresponding fitness inferences are show in Figure S6.

Unlike the OV-075 patient data set, in which the WGD$^+$ population constitutes a majority of the tumor population (WGD$^+$: 96.5%), the OV-006 (WGD$^+$: 7.6%) and OV-139 (WGD$^+$: 12.9%) datasets exhibit substantially smaller WGD$^+$ populations. However, population proportion alone does not directly determine inferred fitness in our framework. Consistent with the primary analysis presented in the main Results section, our LeafRank inference did not support a fitness advantage for WGD$^+$ populations relative to their WGD$^-$ counterparts. Instead, the inferred fitness of WGD$^+$ populations remained comparable to, or lower than, that of their WGD$^-$ sibling populations, suggesting that WGD does not immediately confer a selective advantage.

Similar to the OV-075 analysis in Results, reconstructing the ultrametric trees for subclonal WGD patients remains challenging. In the rootogram, either the WGD$^-$ or WGD$^+$ branches may exhibit systematic deviations that are not fully captured by the current model fit, depending on the patient dataset. One possible explanation is the use of the extreme-case *Anc. assumption*, under which the timing of WGD is placed at the ancestral end of every branch leading to a WGD$^+$ descendant. In addition, technical limitations, including the limited detectability of small-magnitude copy number aberrations and the increased complexity of likelihood optimization under the WGD state-dependent model, may further impede accurate reconstruction of the underlying temporal structure of the ultrametric tree. Improved temporal resolution, particularly regarding the emergence time of WGD$^+$ populations, could substantially improve ultrametric tree reconstruction and downstream fitness inference.

### Robustness of OV-075 data analysis under alternative model assumptions

In the main Results section, our analysis relies on several assumptions regarding the timing of WGD events and the accumulation of copy number aberrations thereafter:

- *Anc. assumption:* For each WGD$^+$ clade, the WGD event is assigned to the ancestral end of the branch leading to the earliest node inferred to be WGD$^+$. Consequently, the branch leading to this earliest WGD$^+$ node is labeled as WGD$^+$, together with all branches descending from this node.

- *Poi. assumption:* The WGD$^+$ population accumulates SCNAs according to the same Poisson process as the WGD$^-$ population, but with an approximately twofold higher aberration rate.

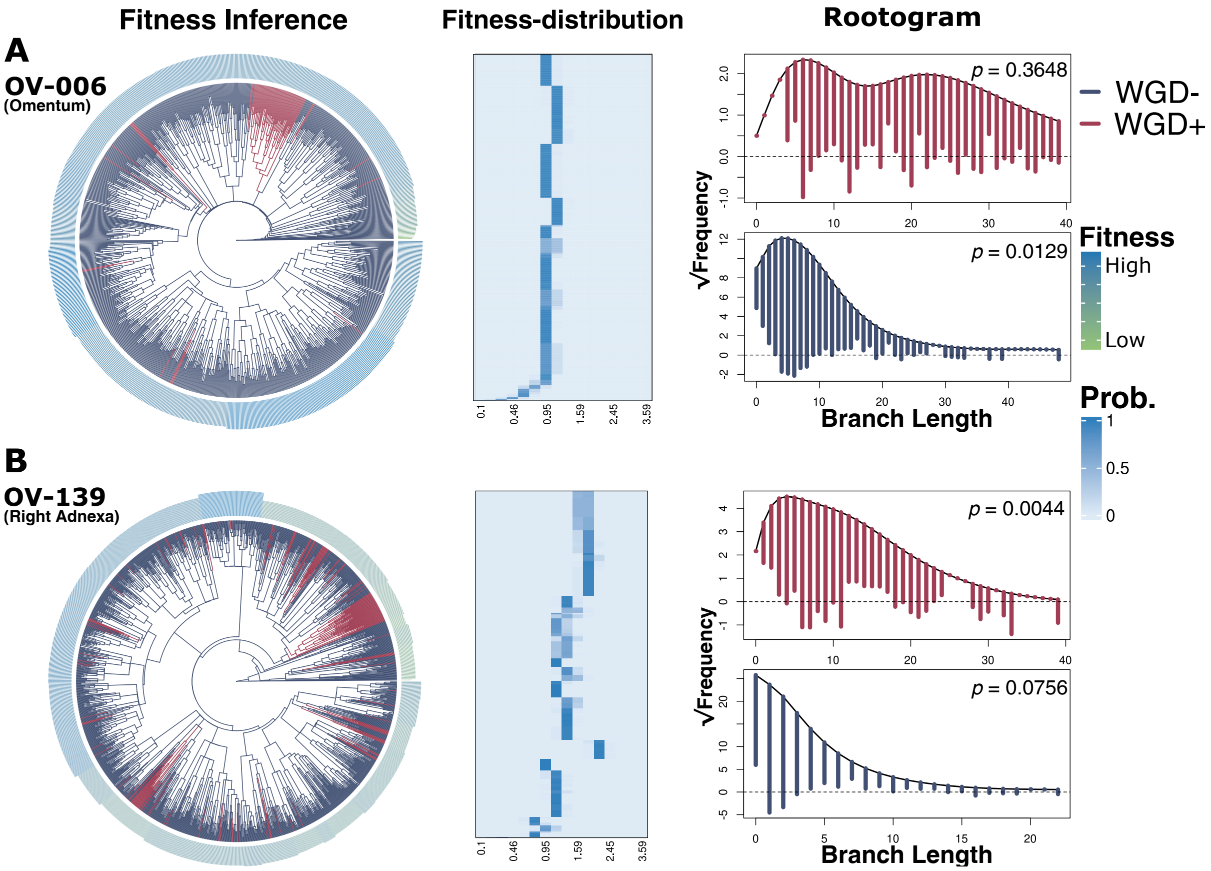


**Figure S6. Subclonal WGD patient data suggest comparable fitness between WGD⁻ and WGD⁺ subclones under a heterogeneous fitness landscape. (A)** Inference results for patient OV-006 with sampling site indicated. The Fitness Inference and Rootogram are presented in the same format as in Figure 7. The Fitness Distribution subpanel presents a heatmap of the marginal distribution of cells across fitness phenotypes. **(B)** Inference results for patient OV-139 with sampling site, shown in the same format as in panel (A).

Here, we evaluate the robustness of our findings under alternative modeling assumptions. Specifically, we consider the following alternatives:

- *Des. assumption:* For each WGD$^+$ clade, the WGD event is assigned to the descendant end of the branch leading to the earliest node inferred to be WGD$^+$. Consequently, branches descending from this earliest WGD $^+$ node are labeled as WGD$^+$, whereas the branch leading to this node is treated as WGD$^-$.
- *N.b. assumption:* The WGD$^+$ population follows the same binomial model for copy number aberration accumulation as the WGD$^+$ population, but with an approximately twofold higher aberration probability.

We first investigated the *Des. assumption* as an alternative framework for branch-level WGD classification. Compared with the *Anc. assumption*, the rootogram in Figure S7B shows no clear improvement in ultrametric tree reconstruction relative to that obtained under the *Anc. assumption*. In addition, this alternative yielded a branch-level WGD classification that was less concordant with the classification reported in the original study[44]. More importantly, reconstruction under the *Des. assumption* implied an approximately 5-fold increase in the copy number aberrations rate in the WGD$^+$ population relative to the WGD$^-$ population. This estimate appears biologically implausible given existing estimates of WGD-associated chromosomal instability[40,44].

Unlike the original *Anc. assumption*, the *Des. assumption* yields a more conservative estimate of WGD timing, thereby shortening the inferred developmental history of the WGD$^+$ population. In practice, the true WGD event must occur somewhere along the branch leading to the earliest node inferred to be WGD$^+$. Consequently, reconstruction under the *Des. assumption* tends to assign higher inferred fitness to WGD$^+$ populations than reconstruction under the *Anc. assumption*.

Indeed, Figure S7A shows that, under the *Des. assumption*, the majority of WGD$^+$ cells exhibit higher inferred fitness than many cells in the WGD$^-$ population. However, as illustrated in the zoom-in panel, a long-persisting, slowly growing WGD$^+$ subpopulation exhibits lower inferred fitness than the high-fitness subgroup within the WGD$^-$ population. Notably, this slow-growing WGD$^+$ lineage shares the same ancestral node as the dominant WGD$^+$ population, suggesting substantial fitness heterogeneity within the WGD$^+$ lineage itself. Therefore, even under the *Des. assumption*, the inferred fitness landscape does not support a consistent or dominant fitness advantage for WGD$^+$ populations.

We next evaluated the alternative *N.b. assumption* for modeling SCNAs. Under this assumption, the model failed to adequately capture the excess of WGD$^+$ branches containing approximately 20 SCNAs. This observation suggests that the Poisson model provided a better fit for the SCNAs distribution in the datasets analyzed here.

Overall, these alternative assumption did not provide a better fit to the data than the assumptions used in the main Results and Methods sections. Under the *Des. assumption*, we found that the primary conclusion regarding the lack of a consistent fitness advantage for WGD$^+$ populations remained qualitatively unchanged. Together, these results support the robustness of the findings presented in the main Results section.

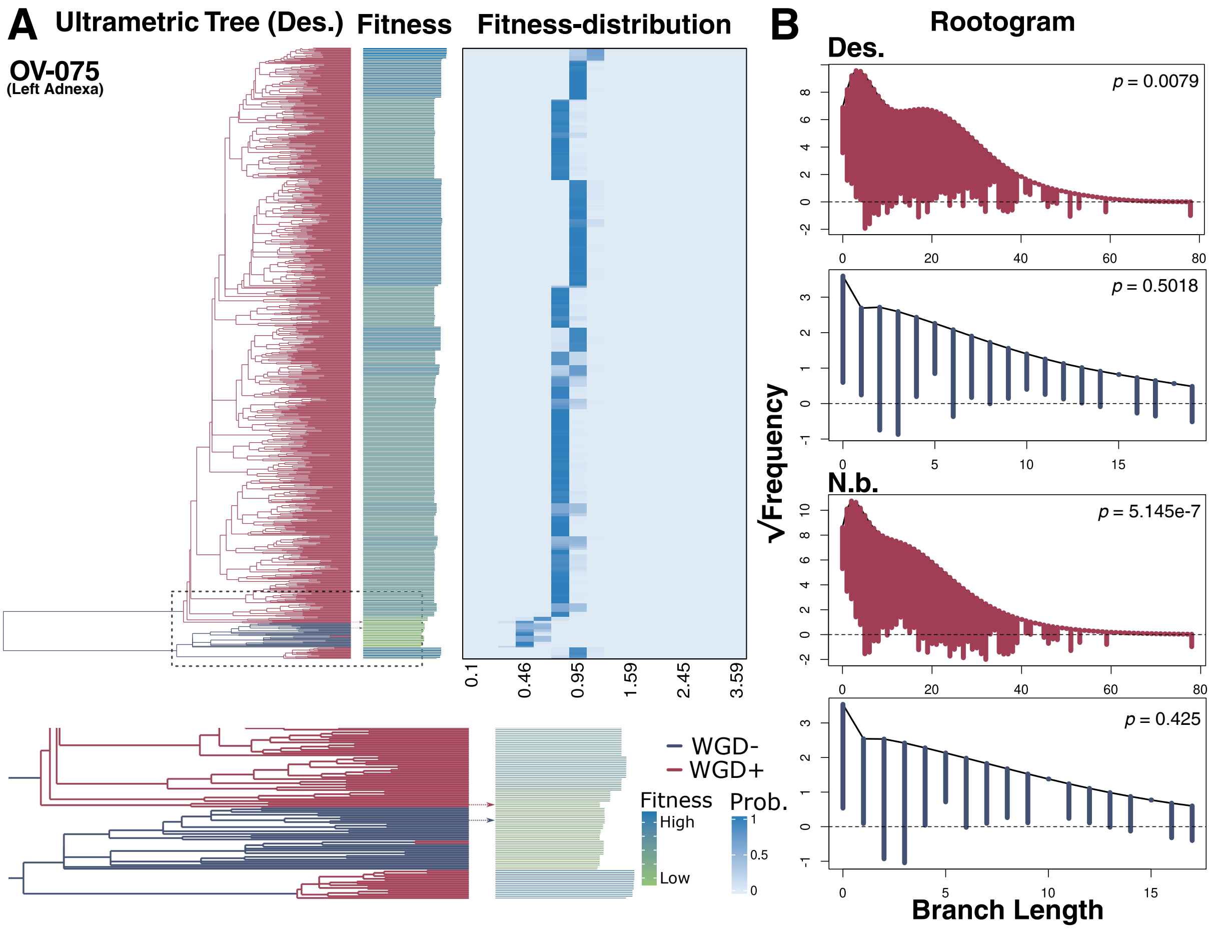


**Figure S7. Fitness inference for OV-075 under alternative assumptions (*Des.* and *N.b.*) yields consistent conclusions regarding the fitness advantage of WGD$^+$ subclones (A)** Inferred fitness results for OV-075 under the *Des. assumption*. The ultrametric tree and fitness distribution subpanels are shown as in Figure S3, with branches color-coded according to WGD status. In the zoom-in panel, inferred fitness values for cells belonging to WGD$^+$ and WGD$^-$ subclones are highlighted. **(B)** Rootograms for ultrametric tree reconstruction under the two alternative assumptions. Results for WGD$^+$ and WGD$^-$ branches are color-coded as in panel A.
